# Requirement for Type IX secretion machinery in a nematode model of Chryseobacterial pathogenesis

**DOI:** 10.64898/2026.09.25.754504

**Authors:** Dae-Eun Jeong, Andrew Z. Fire

## Abstract

Bacterium-nematode interactions in soil environments hold agricultural, economic, and evolutionary significance, with the rapid killing of nematodes by nematophagous bacteria representing an extreme and potentially valuable example of such interactions. Here, we investigate rapid killing of the model nematode *Caenorhabditis elegans* by the nematophagous bacterial species *Chryseobacterium nematophagum* through development and application of bacterial genetic tools for *Chryseobacterium*. We show that *Chryseobacterium nematophagum* can be genetically manipulated through both forward genetics and homology-mediated genetic interventions, with the latter driven by the use of a pyrimidine auxotrophy system to circumvent the multidrug resistance of these bacteria. Combining these approaches, we identified the Type IX Secretion System (T9SS) machinery as a central virulence determinant. Independent mutations in multiple T9SS components each substantially attenuated nematode killing with complementation restoring virulence. This system provides a tractable experimental platform for investigating the molecular basis of nematophagous bacterial pathogenesis and of the T9SS-dependent virulence mechanisms.

## Introduction

Bacteria and their eukaryotic hosts have evolved together through two billion years of interactions ranging from cooperation to predation to pathogenesis [1–4]. All these interactions have shaped the evolution and physiology of both interacting partners [1–6]. Bacteria are particularly adept as pathogens, with the capacity to colonize and exploit hosts across nearly all domains of life, imposing a substantial and ongoing burden through infectious disease, crop loss, and disruption of soil and ecosystem health [7–11].

The genus *Chryseobacterium*, a member of the phylum *Bacteroidota*, comprises a diverse group of Gram-negative, aerobic bacteria distributed broadly across environmental and host-associated niches [12–14]. Members of this genus have been isolated from soil, freshwater, plants, insects, fish, and clinical samples, often displaying strong ecological associations with specific substrates or animal hosts [14]. This breadth of associations is accompanied by functional diversity: *Chryseobacterium* lifestyles range from opportunistic pathogens of fish [15,16] and human [17,18] to colonizers of plant rhizosphere [19–21] and animal microbiota [22–26]. While the environmental impacts of many species of this genus remain unexplored, a few have been shown to contribute to organic matter turnover, including degradation of keratin-rich feathers [27,28] and other protein-rich substrates [14,29–31].

*Chryseobacterium nematophagum* represents a distinct ecological specialization within the genus through its antagonistic interaction with nematodes [32,33]. First isolated and characterized by Felix and Duveau [32], and Page et al. [33], *C. nematophagum* exhibits potent nematophagous activity, including degradation of the chitinous lining of the pharyngeal lumen, colonization of the body cavity, and progressive consumption of internal tissues, a pathogenic strategy that is effective against a broad range of free-living and parasitic nematodes [33]. Unlike the enzymatic degradation of complex environmental substrates that characterizes many saprophytic *Chryseobacterium* species, this nematophagous lifestyle suggests the evolution of specialized virulence strategies specifically directed against a nematode host.

Nematodes are among the most abundant metazoans in terrestrial ecosystems, playing central roles in shaping microbial communities and nutrient cycling [34,35]. The broad nematophagous effects of *C. nematophagum* across free-living and parasitic nematode species positions it as both a promising biocontrol agent and a compelling model for investigating the molecular basis of bacterium-nematode interactions [33]. Comparative genomic analysis by Page et al. [33] identified numerous candidate mechanisms potentially responsible for pathogenicity.

Progress in dissecting the molecular mechanisms underlying nematophagous activity has been limited by the absence of genetic tools for *C. nematophagum* and related *Chryseobacterium* species. Establishing *C. nematophagum* as a genetically tractable model, in conjunction with a well-studied and highly manipulable nematode host *Caenorhabditis elegans* [36–40], offers a unique opportunity to move beyond correlative predictions toward experimental dissection of bacterial mechanisms underlying nematophagous virulence.

Here we describe development of the first genetic manipulation system for *C. nematophagum*, encompassing auxotrophy-based selection, conjugation-mediated plasmid delivery, and targeted gene disruption and complementation, all tools that have been absent for this organism and in the *Chryseobacterium* genus. Leveraging this system with forward genetics, we demonstrate that the Type IX Secretion System (T9SS) machinery constitute a central virulence determinant of *C. nematophagum* against *C. elegans*, as disruption of T9SS components abolishes nematode killing activity. Together, these findings establish *C. nematophagum* as a genetically tractable model organism for studying *Bacteroidota* virulence mechanisms and interkingdom bacterial-nematode interactions, while also laying the groundwork for systematic molecular dissection of nematophagous pathogenesis.

## Results

### Isolation and classification of a nematophagous *Chryseobacterium nematophagum* strain (PDb101) from North America

From a survey aimed at identifying nematode-associating pathogens from soil samples collected in Palo Alto, California, USA, we isolated a bacterial strain exhibiting strong toxicity to nematode *Caenorhabditis elegans*. When *C. elegans* was exposed to this isolate, which we designated PDb101, it exhibited rapid and reproducible mortality compared to animals maintained on *Escherichia coli* (OP50), the standard laboratory food source for *C. elegans* (Fig 1A). *C. elegans* survival was dramatically extended on heat-killed PDb101 relative to live bacteria (Fig 1B), demonstrating that pathogenic activity requires live bacteria rather than pre-formed heat-stable factors. In addition, treatment of PDb101 with 5-fluoro-2′-deoxyuridine (FUdR) that kills proliferating PDb101 (Fig S1A-B) reduced *C. elegans* killing in a dose-dependent manner (Fig S1C), suggesting that active bacterial proliferation contributes to pathogenicity.

**Fig 1.**
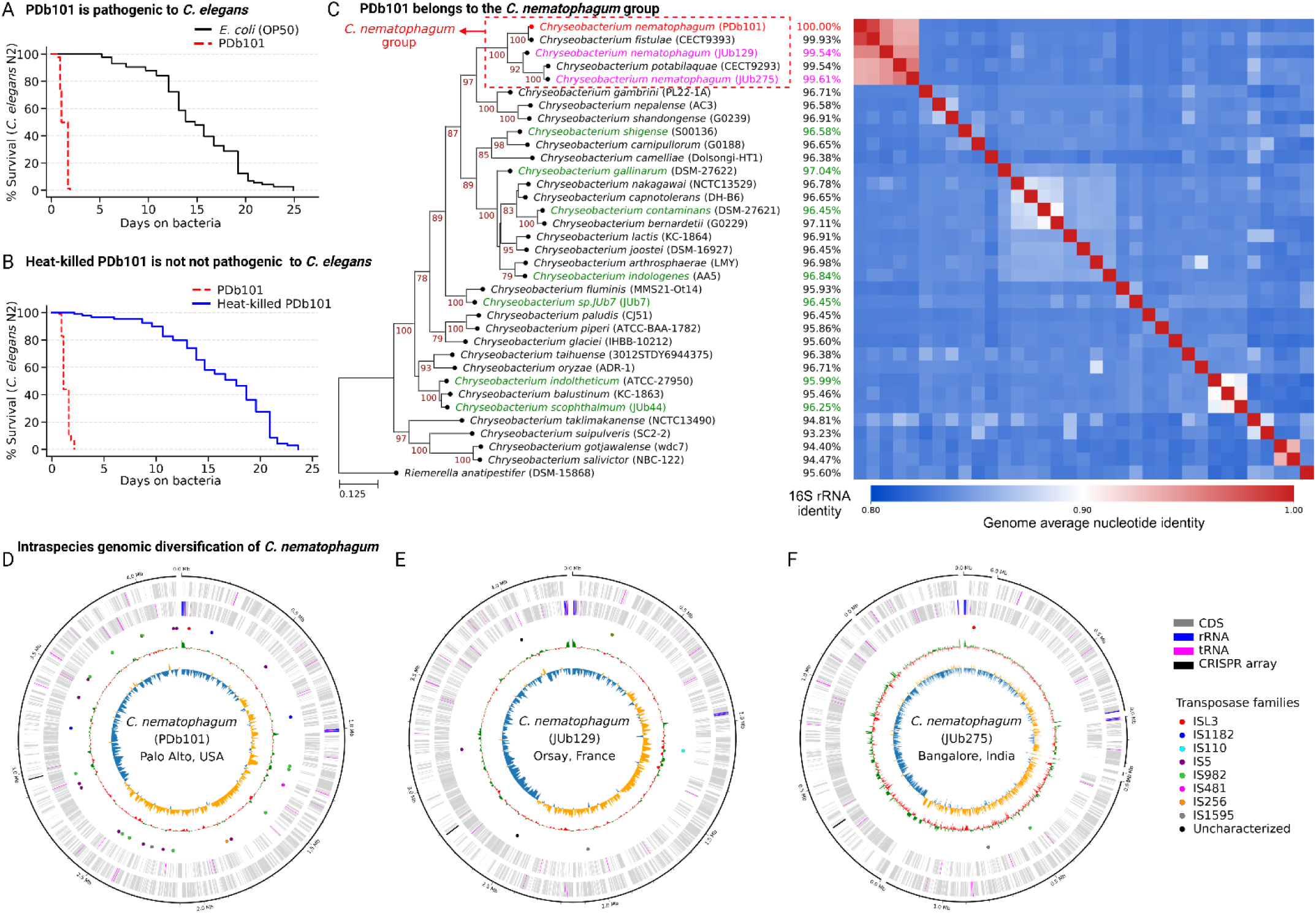
PDb101 is *C. nematophagum*, pathogenic bacteria to nematode *C. elegans*. (**A**) Survival curves of nematode *C. elegans* (N2) exposed to bacteria *E. coli* (OP50) or PDb101. (**B**) Survival curves of *C. elegans* (N2) on live PDb101 or heat-killed PDb101. See Supplementary Table S4 for additional repeats and statistical analysis for the survival data. (**C**) A pan-genome scale phylogenetic tree of bacterial species in the *Chryseobacterium* genus (Supplementary Table S2) was constructed with a maximum likelihood (ML) method (IQ-TREE) based on a multiple sequence alignment of conserved proteins from the genomes (Roary). The numbers on the top position of tree branches are bootstrap supporting values. Red (PDb101) and magenta (JUb129 and JUb275) colors denote pathogenic *C. nematophagum* species. Green color indicates non-pathogenic *Chryseobacterium* species shown in a previous study [33]. The first right column of the tree indicates 16S rRNA sequence identities to that of PDb101. A heat map shows pairwise comparison of genome average nucleotide identities to the PDb101 genome (pyANI-plus). (**D-F**) Circular genome maps (Circos) of PDb101 (D), JUb129 (E) and JUb275 (F) *C. nematophagum*. From the edge of the circle into the center, the first layer shows contig length. The second and third layers show genes with forward and reverse directions with colors; gray for CDS, blue for rRNA, magenta for tRNA, and black for CRISPR array. In the fourth layer, the small circles and their respective colors indicate the position and the families of transposase enzymes found in the genomes (ISEScan). The fifth and sixth layers show GC content and GC skew of the genomes.

To determine the taxonomic identity of PDb101, we sequenced the whole genome and performed phylogenetic analyses. Sequence-based classification revealed that PDb101 belongs to the genus *Chryseobacterium*. Analysis of the 16S rRNA genes and genome-wide comparisons showed that PDb101 shares highly similar sequences with previously described *Chryseobacterium* species isolated from geographically distant locations, including *C. fistulae* and *C. potabilaquae* (Spain) [41], and two *C. nematophagum* strains, JUb129 (France) and JUb275 (India) [33] (Fig 1C).

Consistent with the nematopathogenic activity previously reported for two *C. nematophagum* strains [33], rapid pharyngeal destruction was visible after 3 hours of exposure to PDb101 (Fig S1D). A pharyngeal muscle reporter *C. elegans* (*myo2-p::mCherry*) showed distortion of the pharyngeal axis upon exposure to PDb101 while animals fed *E. coli* OP50 maintained intact pharyngeal structure (Fig S1D). Based on these pathogenic activities in conjunction with phylogenetic analyses, PDb101 was assigned to the *C. nematophagum* group (red-dashed box in Fig 1C).

**Fig S1.**
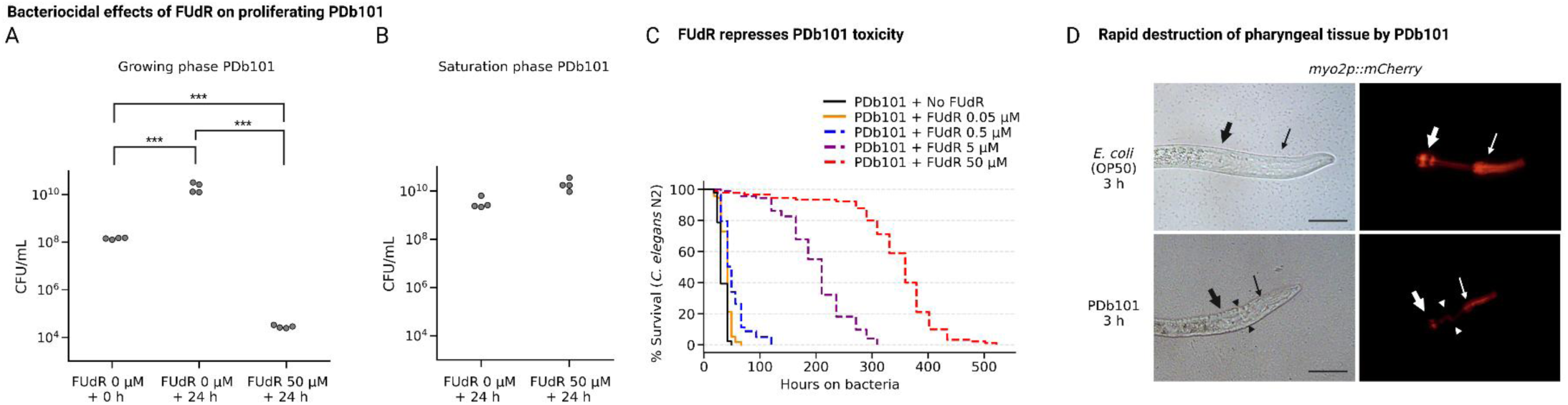
Characterization of FUdR effect on PDb101 toxicity and *C. elegans* pharyngeal tissue damage by PDb101. (**A**) Categorical scatter plots showing colony forming units per mL (CFU/mL) of PDb101 cultures under three conditions: untreated cells at time zero (FUdR 0 µM + 0 h), untreated cells after 24 hours of incubation in 1X PGM media (FUdR 0 µM + 24 h), and cells treated with 50 µM FUdR after 24 hours of incubation in 1X PGM media (FUdR 50 µM + 24 h). FUdR treatment at 50 µM significantly reduced viable cell counts relative to both the untreated 24-hour control and the time-zero baseline (*** *p* < 0.001, one-way ANOVA-Tukey HSD post-hoc test), confirming that FUdR exerts bactericidal activity against PDb101. (**B**) CFU/mL of saturation phase PDb101 cultures in the absence (FUdR 0 µM + 24 h) or presence (FUdR 50 µM + 24 h) of 50 µM FUdR after 24 hours of incubation. In contrast to growing phase cells, saturation phase PDb101 cells were resistant to FUdR treatment, with no significant reduction in viable cell counts observed, indicating that FUdR bactericidal activity is selective for proliferating cells. (**C**) Kaplan-Meier survival curves of *C. elegans* N2 adult animals fed on PDb101 in the absence or presence of increasing concentrations of FUdR (0.05, 0.5, 5, and 50 µM). FUdR suppressed PDb101-dependent nematode killing in a dose-dependent manner, with survival progressively extended at higher FUdR concentrations. (**D**) Brightfield (left) and fluorescence (right) micrographs of *C. elegans* L1 stage animals expressing a pharyngeal tissue reporter (VS21, *myo-2p::mCherry*) after 3 hours of exposure to *E. coli* OP50 (top panels) or PDb101 (bottom panels). Notably, while *C. elegans* animals fed with *E. coli* OP50 display a straight linear alignment between the anterior and posterior pharyngeal bulbs (thin and thick arrows, respectively), animals exposed to PDb101 for 3 hours exhibit a distortion of the pharyngeal axis (indicated by arrowheads), suggesting rapid structural disruption of the pharyngeal isthmus by PDb101.

**Fig S2.**
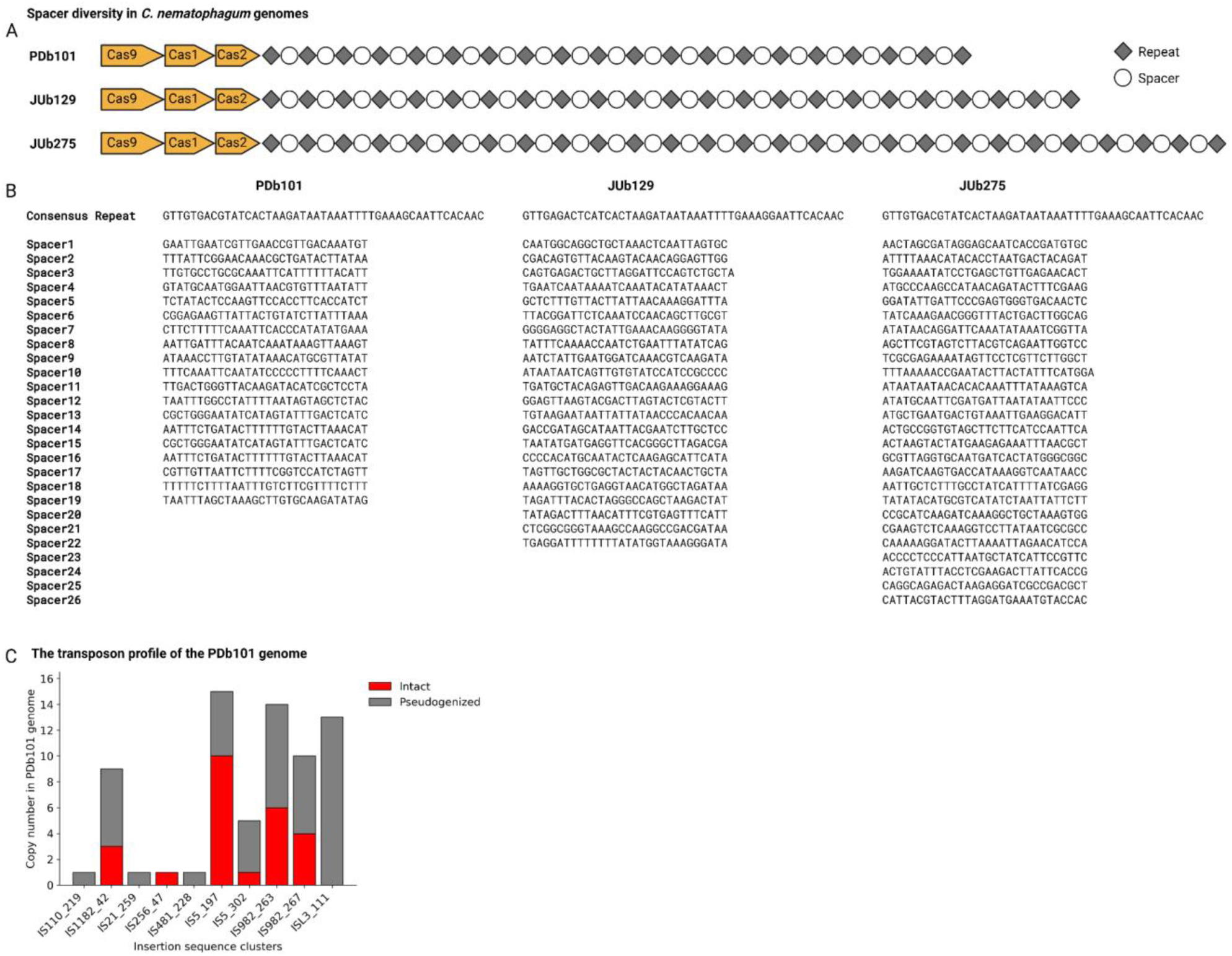
CRISPR spacer diversity and transposon landscape in *Chryseobacterium nematophagum* genomes. (**A-B**) A schematic representation of CRISPR array organization in three *C. nematophagum* strains (PDb101, JUb129 and JUb275) (A). Each array consists of a Cas9-Cas1-Cas2 operon followed by alternating repeat units (gray diamonds) and spacers (empty circles). All repeats are nearly identical, but every spacer has a unique sequence as shown in panel (B), suggesting that three strains had been exposed to different viral or mobile genetic elements, resulting in the acquisition of diverse spacers at sequence and copy number levels. (**C**) A bar chart showing the copy number of insertion sequence (IS) element clusters identified in the PDb101 genome by ISEScan. Each cluster is labeled by IS family designation on the x-axis. Red bars indicate intact IS elements; gray bars indicate pseudogenized copies.

To further examine genomic diversity within the *C. nematophagum* group, we conducted intra-species comparative genomic analyses. This analysis revealed nearly identical CRISPR repeat sequences but substantial variation in CRISPR spacer identity and copy number among strains, consistent with independent exposure to distinct mobile genetic elements and/or phage populations (Fig S2A-B). In addition, both the distribution and diversity of transposable elements varied markedly across strains (Fig 1D-F). Notably, PDb101 harbored a larger number of transposons that were broadly distributed throughout the genome compared to other *C. nematophagum* isolates (Fig 1D-F & Fig S2C). This extensive transposon dispersion suggests that PDb101 has experienced repeated or prolonged exposure to diverse mobile genetic elements during its evolutionary history.

The identification of PDb101 extends the known geographic distribution of nematophagous *C. nematophagum* group beyond Europe and Asia to the American continent, establishing PDb101 as the first North American isolate of this species. This finding is consistent with the global distribution of host nematodes including *C. elegans* and suggests that interactions between nematodes and nematophagous *Chryseobacterium* species may be widespread across terrestrial ecosystems rather than restricted to specific geographic regions.

**Fig S3.**
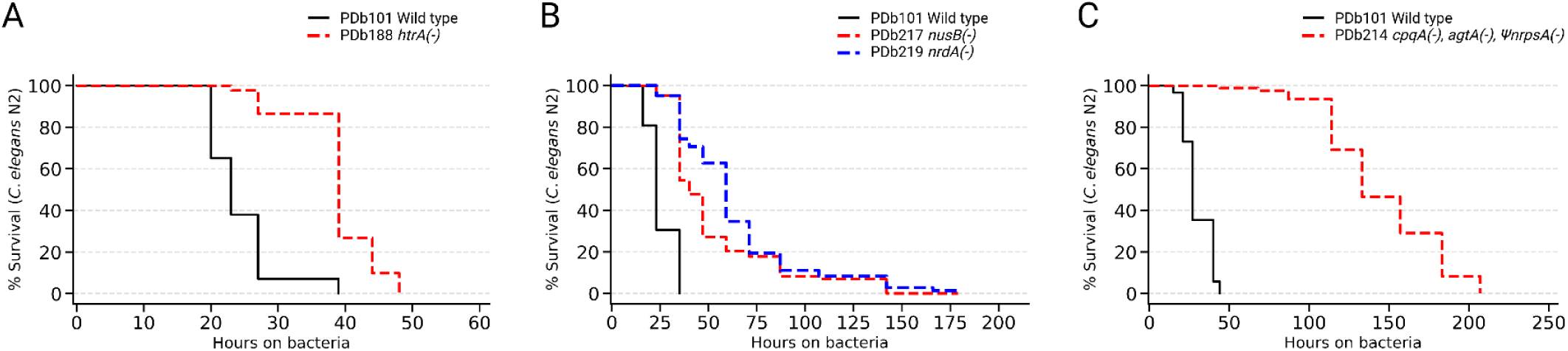
EMS random mutagenesis identified *C. nematophagum* mutants with reduced toxicity to *C. elegans*. (**A–C**) Kaplan-Meier survival curves of *C. elegans* N2 animals fed on wild-type PDb101 (black) or the indicated EMS-induced mutant strains (red or blue dashed), recovered from independent EMS mutagenesis experiments. **(A)** PDb188 carries a G136R missense substitution in *htrA*, encoding a TetR family HTH-type transcriptional regulator, and exhibits moderately reduced killing kinetics relative to wild-type PDb101. (**B**) PDb217 carries a G289D missense substitution in *nusB*, encoding transcription antitermination factor NusB, and PDb219 carries an R385Q missense substitution in *nrdA*, encoding ribonucleoside-diphosphate reductase subunit alpha; both mutants display mild to moderate virulence attenuation relative to wild-type PDb101. (**C**) PDb214 exhibits substantially attenuated killing activity and carries three EMS-induced mutations: a W309Stop nonsense mutation in *cpqA* encoding carboxypeptidase Q, a G725E missense substitution in *agtA* encoding 4-alpha-N-acetylgalactosaminyltransferase, and an E20K missense substitution in a pseudogene *ΨnrpsA* encoding non-ribosomal-peptide-synthase (NRPS); the causative allele responsible for virulence attenuation has not yet been determined. See Supplementary Table S4 for statistical analysis for the survival data.

### EMS mutagenesis identifies genes contributing to virulence

As a first attempt to understand nematophagous activity at the genetic level, we used ethyl methanesulfonate (EMS) to mutagenize PDb101. After optimizing EMS treatment conditions (Supplementary Table S3, and see Materials and Methods), we performed EMS random mutagenesis to generate a population of PDb101 mutant strains, which we then subjected to testing nematophagous activity. The vast majority of the mutant strains (over 4,000 strains) retained intact pathogenicity, with only four mutant strains showing compromised killing (Fig S3A-C). For three of these strains the attenuation was only modest (the strains remained somewhat toxic to *C. elegans*, Fig S3A-B), while a fourth strain had more substantial attenuation of killing but was found to carry three separate mutations (Fig S3C, see the figure legend for details). The multiple mutations in the virulence-attenuated strain highlight a key limitation of EMS mutagenesis. Its random and genome-wide nature makes it difficult to link phenotype to a single causal mutation. Robust dissection of PDb101 virulence therefore requires complementation analysis and targeted genetic tools, such as directed mutagenesis approaches.

### Establishment of a genetic manipulation system in *Chryseobacterium nematophagum* (PDb101)

Genetic dissection of the strong nematode-killing activity exhibited by PDb101 would require a reliable genetic manipulation system. As an initial approach, we considered selecting transformed PDb101 cells using antibiotic resistance markers. However, PDb101 exhibited intrinsic resistance to a broad range of commonly used antibiotics, including ampicillin (beta-lactam), kanamycin/gentamicin (aminoglycoside), spectinomycin (aminocyclitol), erythromycin (macrolides), chloramphenicol, tetracycline and trimethoprim, rendering these markers ineffective for selection. Although PDb101 was sensitive to vancomycin (glycopeptide) and rifampicin (ansamycin), resistance to these antibiotics was not pursued as a selection strategy due to antibiotic stewardship and their limited suitability for routine bacterial genetic manipulation.

To develop a safer and more robust selection system, we instead explored the use of auxotrophic markers for selection systems. PDb101 grew robustly on complex media supplemented with peptone or yeast extract/tryptone, prompting us to identify chemically defined minimal media conditions that support its growth. We found that PDb101 was capable of growth on M9 minimal medium supplemented with glucose, a defined mixture of 19 amino acids and thiamine (vitamin B1), establishing conditions suitable for metabolic selection.

From the mutagenized population, we isolated two independent mutant strains that grew normally on complex media but failed to grow on minimal media (Fig 2A). Genome analysis of these mutants revealed mutations affecting two distinct genes in the pyrimidine biosynthesis pathway (Fig 2B). One mutant carried a mutation in the gene encoding dihydroorotase (*DHO*), whereas the second mutant harbored a mutation in the gene encoding the carbamoyl-phosphate synthase (*CPS*) large subunit. Both mutations are missense substitutions affecting residues that are conserved across bacteria and eukaryotes: a G53R substitution in DHO and an E755K substitution in the CPS large subunit (Fig S4A-B). The conservation of these residues across phylogenetically diverse organisms, including animals, suggests their roles in catalysis and/or structural integrity, and explains the auxotrophic phenotype conferred by each mutation. Importantly, supplementation of minimal media with exogenous uracil fully restored growth of both the *DHO* and *CPS* mutants (Fig 2C-E), confirming that the growth defect in each strain results specifically from impaired *de novo* pyrimidine biosynthesis rather than pleiotropic effects of the EMS treatment. These auxotrophic strains therefore provide a clean metabolic selection system in which growth on minimal media serves as a direct readout of functional complementation. The *DHO* gene, due to its relatively small size, non-operonic genomic context and its little impact on the host virulence on complex media (Fig S4C), was selected as a potential complementation marker for subsequent genetic manipulation experiments.

**Fig 2.**
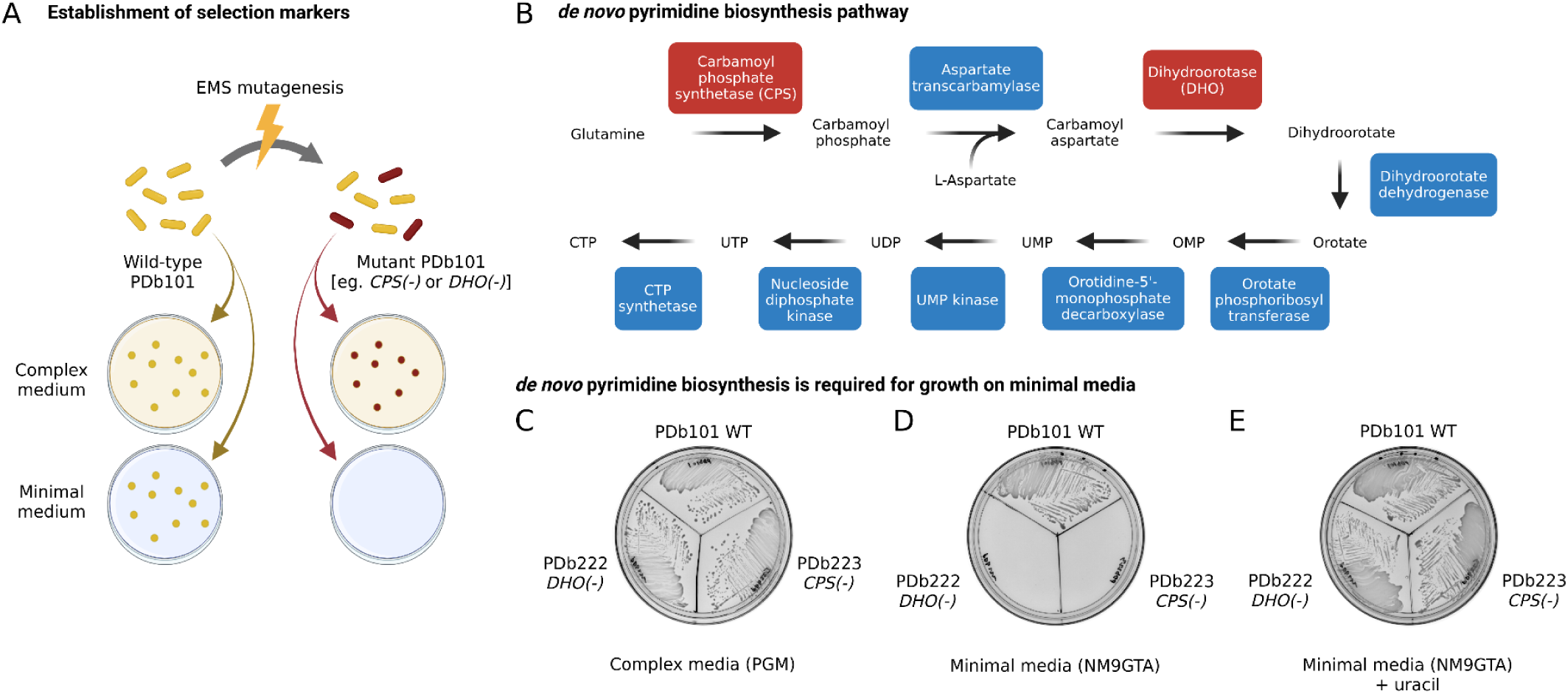
*de novo* pyrimidine biosynthesis is required for growth of PDb101 under the minimal media condition. (**A**) A schematic of EMS mutagenesis strategy to generate auxotrophic mutants capable of growth on complex media but not on minimal media. (**B**) A schematic of the *de novo* pyrimidine biosynthesis pathway. Enzymatic steps highlighted in red (Carbamoyl phosphate synthetase, CPS; Dihydroorotase, DHO) represent the mutated loci characterized in this study. (**C-E**) Growth phenotypes of wild-type PDb101, PDb222 *DHO(-)*, and PDb223 *CPS(-)* on (C) complex medium (PGM), (D) minimal medium (NM9GTA), and (E) minimal medium supplemented with uracil (NM9GTA + uracil).

**Fig S4.**
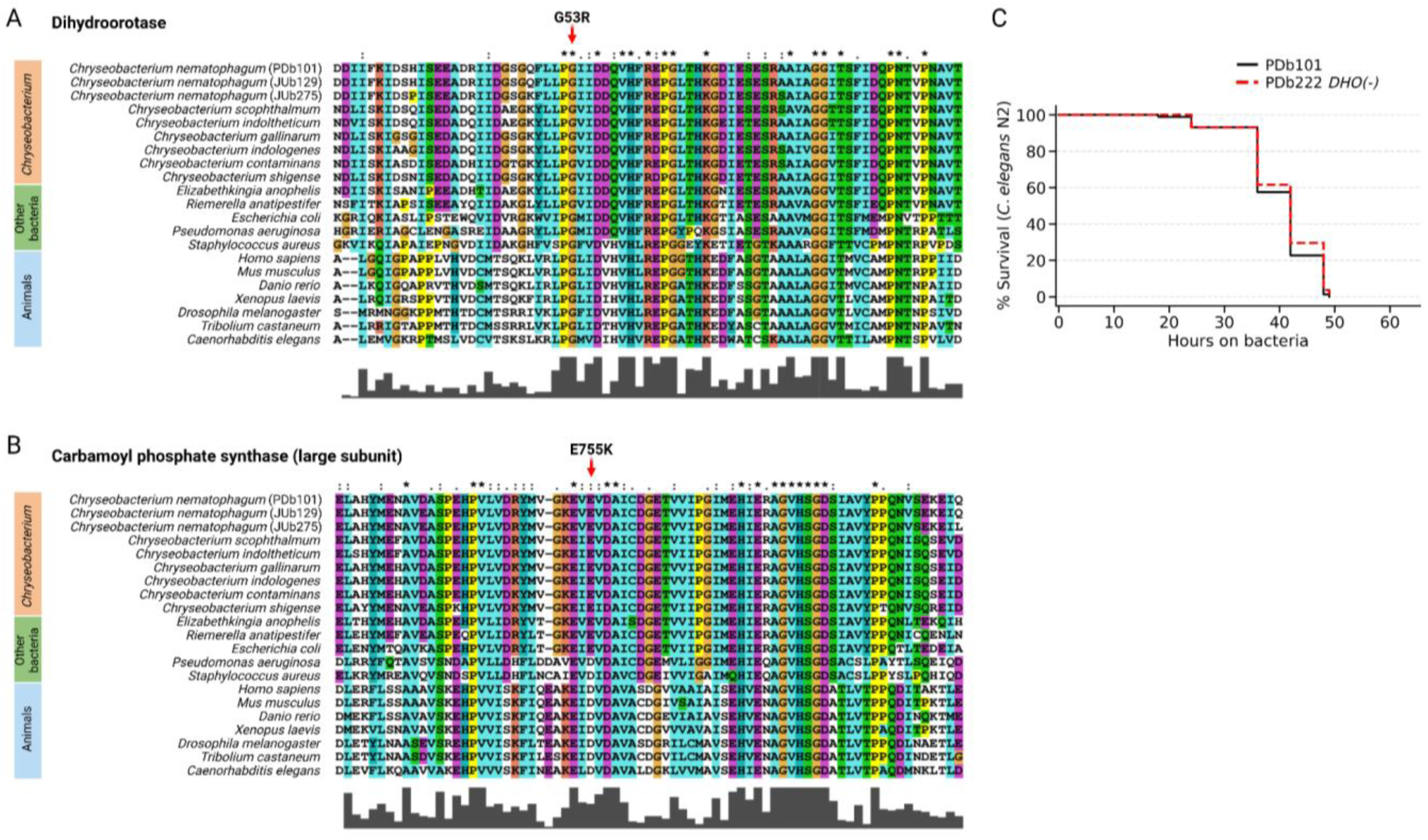
Multiple sequence alignments of dihydroorotase and carbamoyl phosphate synthetase (large subunit), and virulence phenotype of the *DHO(-)* auxotroph. **(A-B)** Multiple sequence alignment of Dihydroorotase (DHO) (A) and carbamoyl phosphate synthetase large subunit (CPS) (B) protein sequences from *Chryseobacterium* genus species, other genus bacteria, and animals. The EMS-induced missense mutations are indicated by red arrows above the alignment. Asterisks and colons below the alignment denote fully conserved and biochemically similar positions, respectively; bar height indicates per-column conservation score. Both G53 and E755, for DHO and CPS respectively, fall within a conserved region shared across bacterial and animal protein sequences, consistent with a functionally deleterious substitution. **(C)** Kaplan-Meier survival curves of *C. elegans* strain N2 (CGC1) fed on wild-type PDb101 (black circles) or the *DHO(-)* auxotroph PDb222 (red squares). Both strains kill *C. elegans* with similar kinetics over approximately 48 hours, indicating that pyrimidine auxotrophy does not attenuate the virulence of PDb101 under the standard *C. elegans* killing assay conditions.

To test whether the *DHO* gene could serve as a selectable marker for genetic manipulation of PDb101, the wild-type *DHO* coding sequence was cloned into a broad-host-range plasmid and introduced into an *E. coli* conjugation donor strain. Delivery of this plasmid into *DHO*-deficient PDb101 (PDb222) by conjugation (Fig 3A) restored the ability of transconjugant cells to grow on minimal medium, demonstrating successful plasmid transfer and functional complementation of the auxotrophy (Fig 3B-D). Interestingly, genome sequencing of transconjugant isolates revealed that the plasmid had integrated into the chromosome as a whole-plasmid insertion rather than being maintained as an extrachromosomal element. The integrated plasmid was found at the mutant *DHO* locus, indicating that recombination had occurred via sequence homology between the plasmid-borne *DHO^WT^* and the chromosomal *DHO^Mut^* allele (Fig S5A). It is likely that chromosomal integrants were selectively enriched over plasmid-bearing cells during serial passaging on both liquid and solid minimal media, likely due to a growth advantage conferred by stable chromosomal inheritance of the *DHO^WT^* marker over low copy plasmid maintenance. Importantly, this recombination-driven whole-plasmid insertion phenomenon can be exploited as a general strategy for targeted chromosomal manipulation: by providing a single homology arm corresponding to any genomic locus of interest on the plasmid, site-specific integration and targeted disruption of that locus can be achieved (Fig S5B).

**Fig 3.**
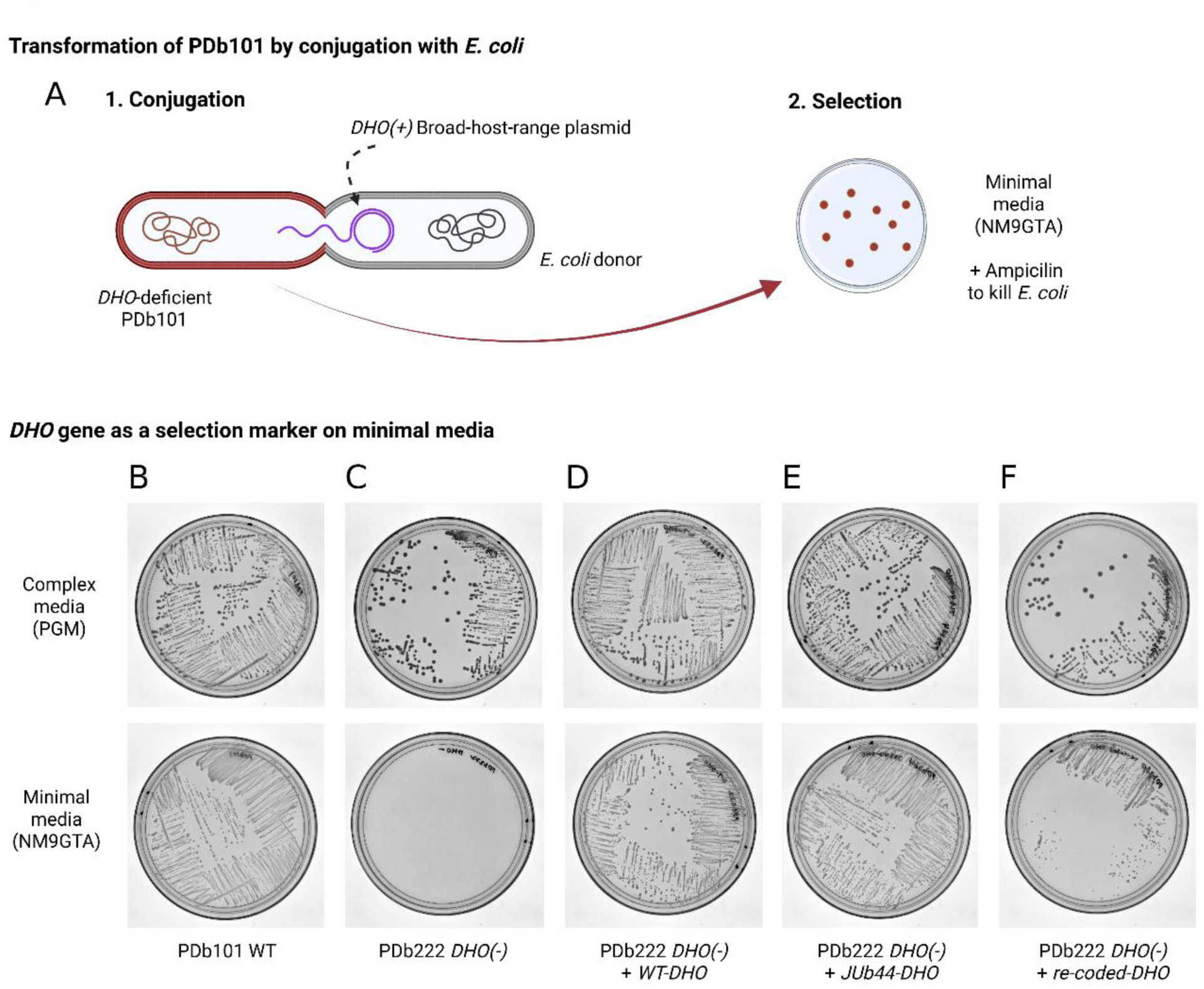
Conjugation-mediated delivery of a *DHO* gene restores growth of *DHO*-deficient PDb101 on minimal media. (**A**) A schematic of the conjugation and selection strategy. A broad-host-range plasmid carrying a functional *DHO* gene was transferred from an *E. coli* donor strain into *DHO*-deficient PDb101 (PDb222) by conjugation. Transconjugants were selected on minimal medium (NM9GTA) supplemented with ampicillin to eliminate residual *E. coli* donor cells, with colony growth indicating successful plasmid acquisition and complementation of the pyrimidine auxotrophy. **(B-F)** Growth of the indicated strains on complex medium (PGM; top row) and minimal medium (NM9GTA; bottom row). Wild-type PDb101 grows on both media (B). The *DHO(-)* auxotroph PDb222 grows on complex medium but fails to grow on minimal medium (C). PDb222 complemented with a plasmid carrying the wild-type *DHO* allele (*WT-DHO*) (D), or the *DHO* allele derived from *C. scophthalmum* strain JUb44 (*JU44b-DHO*) (E), or a codon-recoded *DHO* construct (*re-coded-DHO*) (F) restores growth, validating *DHO* as a functional selectable marker for transformation.

To implement this strategy for genes other than *DHO*, it was necessary to prevent unintended recombination between the plasmid-borne *DHO* marker and the endogenous *DHO^Mut^* locus. We therefore tested two alternative *DHO* constructs as selectable markers: the *DHO* gene (JUb44-DHO) from a closely related species, *Chryseobacterium scophthalmum* (JUb44), and a codon-recoded version of the PDb101 *DHO* gene (re-coded-DHO) encoding an identical protein sequence but with altered codon usage. Both constructs successfully restored growth of *DHO*-deficient PDb222 on minimal medium (Fig 3E-F), validating their use as recombination-insulating selectable markers for targeted gene manipulation in PDb101.

**Fig S5.**
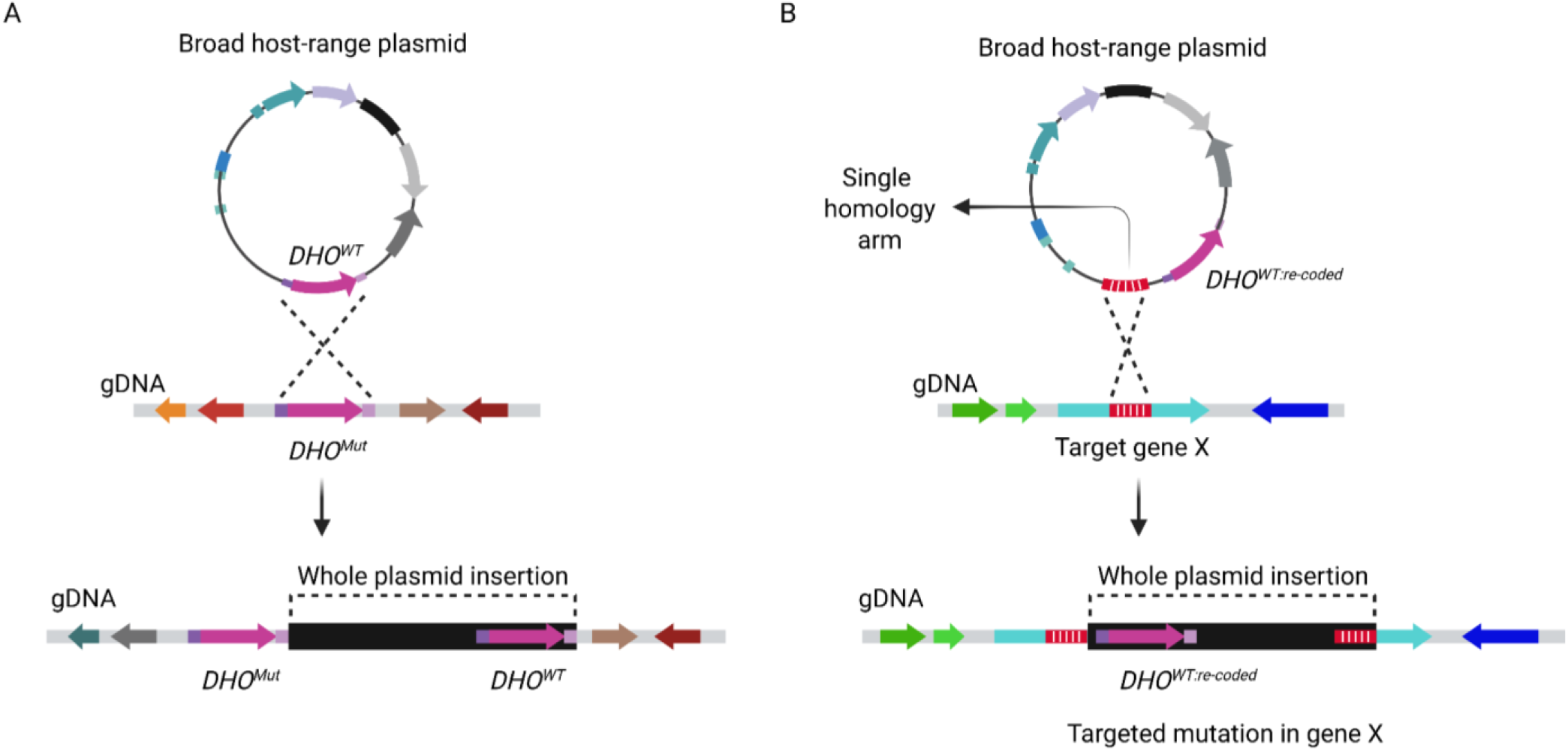
Single homology arm strategy for chromosomal gene targeting in PDb101. (**A**) A schematic of the *DHO* complementation control using whole-plasmid insertion. A broad-host-range plasmid carrying the wild-type *DHO* allele (*DHO^WT^*) shares sequence homology with the mutant *DHO* locus (*DHO^Mut^*) on the PDb222 chromosome. Recombination between the plasmid-borne *DHO*^WT^ and the chromosomal *DHO^Mut^* via a single crossover results in whole-plasmid integration, yielding a chromosomal configuration in which *DHO^Mut^* and *DHO^WT^* flank the inserted plasmid backbone. This arrangement restores a functional *DHO* copy in *cis* while retaining the original mutant allele. (**B**) A schematic of the single homology arm strategy for targeted mutagenesis of an arbitrary gene of interest (Target gene X). A broad-host-range plasmid carries a codon-recoded *DHO* selectable marker (*DHO^WT:re-coded^*) and a single homology arm (red/white hatched segment) corresponding to an internal region of Target gene X. The *DHO* allele is codon-recoded to prevent unintended recombination with the endogenous *DHO^Mut^* locus during targeting of unrelated genomic loci. Recombination between the plasmid-borne homology arm and the matching chromosomal sequence via a single crossover drives whole-plasmid insertion into gene X, simultaneously disrupting the target and integrating the *DHO^WT:re-coded^* selectable marker. The resulting chromosomal configuration contains a split, non-functional copy of gene X flanking the inserted plasmid backbone, and integrants are selected on minimal medium lacking uracil. This strategy generalizes the *DHO*-based selection system for targeted gene disruption at any locus in the PDb101 genome except at essential loci.

### Type IX secretion/gliding motility genes contribute to the PDb101 virulence

Independent of the earlier EMS screen, we serendipitously identified seven isolates displaying significantly attenuated *C. elegans* killing activity. These virulence-impaired strains were recovered during the establishment of molecular genetic toolbox for targeted mutagenesis; however, the precise mechanism underlying their spontaneous emergence remains currently undetermined (considered further in the Discussion below). To determine the genetic basis of this attenuated virulence, we performed long-read genome sequencing on each of these *C. nematophagum* isolates. The sequencing identified seven independent mutations (one in each strain) in a set of genes with defined roles in type IX secretion system (T9SS) machinery (Fig 4A): *sprA*, *gldG*, the *porU/porV* operon, *gldJ*, and *gldK* (Fig 4B–E). The convergence of independent mutations on this specific gene set prompted us to examine the biological functions of these loci more closely.

**Fig 4.**
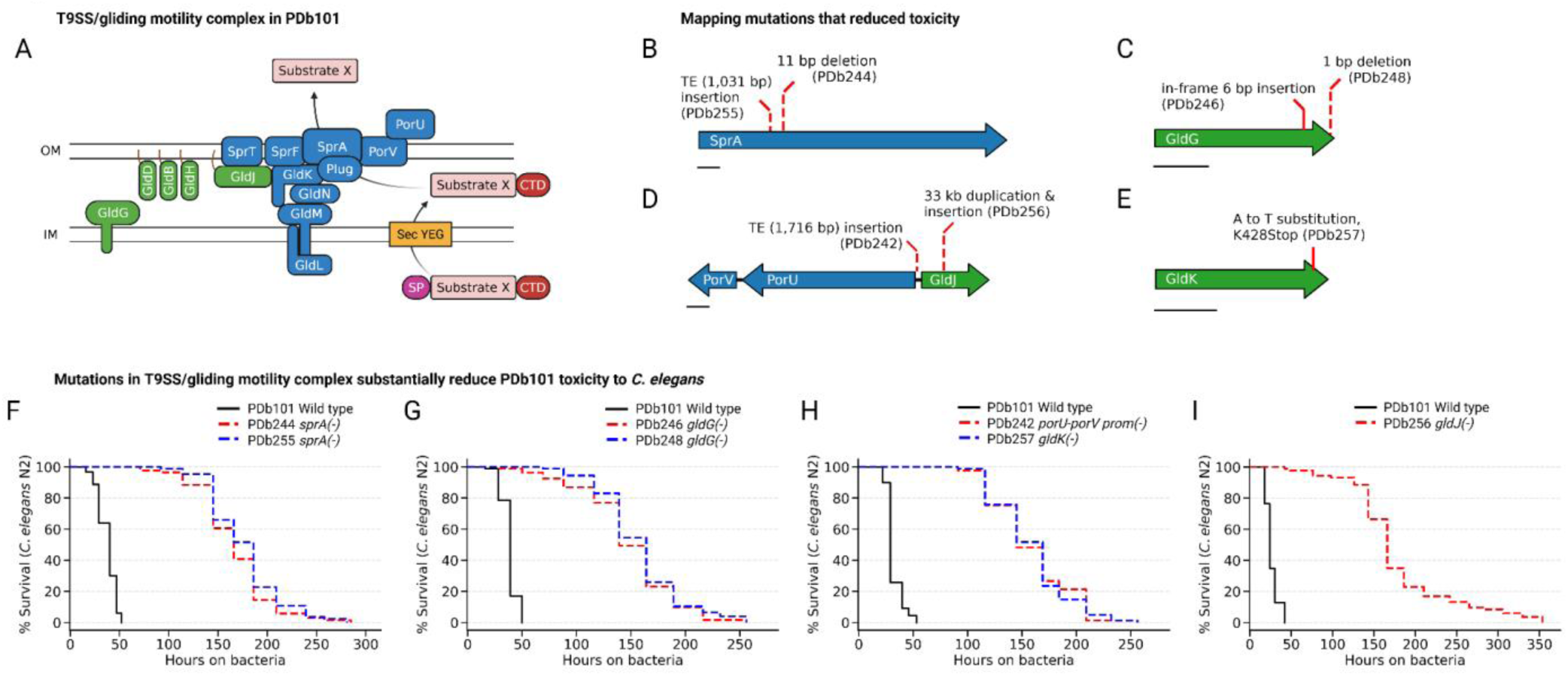
Mutations in the T9SS/gliding motility complex substantially reduce PDb101 toxicity to *C. elegans*. (**A**) A schematic of the T9SS/gliding motility complex in PDb101. The inner membrane (IM) motor module consists of GldL, GldM and GldG, which transduce proton motive force to power both gliding motility and T9SS-dependent secretion. The outer membrane (OM) platform comprises SprA, SprF, SprT, GldJ, GldK, GldN, PorU, and PorV. T9SS cargo substrates bearing a C-terminal domain (CTD) are exported across the outer membrane after Sec-dependent translocation across the inner membrane. (**B–E**) Gene-level schematics illustrating the nature and position of mutations recovered in strains carrying mutations in T9SS/gliding motility complex components. Mutation positions are indicated by red dashed lines above each gene arrow. Scale bars: 500 bp. (**B**) Two independent *sprA* alleles: a transposable element (TE) insertion of 1,031 bp (PDb255) and an 11 bp deletion (PDb244). (**C**) Two independent *gldG* alleles: an in-frame 6 bp insertion (PDb246) and a 1 bp deletion causing frameshift (PDb248). (**D**) Mutations disrupting the *porU-porV-gldJ* locus: a TE insertion of 1,716 bp within the *porU-porV* promoter region (PDb242), and a 33 kb duplication and insertion at the *gldJ* locus (PDb256). (**E**) A nonsense mutation in *gldK*: an A-to-T transversion generating a K428Stop allele (PDb257). (**F–J**) Kaplan-Meier survival curves of *C. elegans* N2 animals fed on wild-type PDb101 (black) or the indicated mutant strains (red dashed, blue dotted). All mutants exhibit substantially delayed killing kinetics relative to wild-type PDb101. (**F**) Two independent *sprA(-)* alleles (PDb244, PDb255). (**G**) Two independent *gldG(-)* alleles (PDb246, PDb248). (**H**) A *porU-porV* promoter disruption allele (PDb242) and a *gldK(-)* allele (PDb257). **(I)** A *gldJ(-)* allele (PDb256). See Supplementary Table S4 for additional repeats and statistical analysis for the survival data.

The Type IX Secretion System (T9SS) is a bacterial secretion apparatus, exclusive to the phylum *Bacteroidota*, that exports cargo proteins bearing a conserved C-terminal domain (CTD) across the outer membrane [42–46]. Substrates are first translocated across the inner membrane via the N-terminal signal peptide (SP)-Sec pathway, then exported through the SprA outer membrane pore, where PorV and PorU/PorZ process the CTD and mediate surface attachment or release of the mature protein [46–52]. The T9SS is physically and functionally coupled to the gliding motility apparatus through the shared inner membrane motor GldL/GldM, with the full complex requiring additional inner membrane (ex. GldG), periplasmic and outer membrane components (ex. GldJ and GldK) for stable assembly and function [42,46,53–57] (Fig 4A). This mechanistic coupling between protein secretion and motility, in which the two processes share the same core protein components, distinguished the T9SS from other known bacterial secretion systems [42,46]. T9SS-dependent secretion and/or gliding motility has been established as a central virulence determinant in the periodontal pathogen *Porphyromonas gingivalis* [42,58–63] and the fish pathogens *Flavobacterium columnare* [64] and *F. psychrophilum* [65,66], but its role in nematophagous activity of *C. nematophagum* has not been experimentally investigated.

Two independent alleles were recovered for *sprA*: PDb255 carries a IS1182 family transposable element insertion (1,031 bp) within the *sprA* coding sequence, and PDb244 carries an 11 bp deletion predicted to cause a frameshift and premature truncation (Fig 4B). SprA is a large outer membrane β-barrel protein that forms the primary translocation pore through which T9SS cargo substrates are exported to the cell surface [46]; both alleles are predicted to compromise pore formation and thereby block substrate secretion. Two independent alleles were likewise recovered for *gldG*: PDb246 carries an in-frame 6 bp insertion, while PDb248 carries a 1 bp deletion causing frameshift at the C-terminus (Fig 4C). GldG is an inner membrane-anchored component of the T9SS/gliding motility complex that contains an ABC transporter-like domain and plays a structural role in linking the inner membrane motor to the outer membrane secretion apparatus [46,67]; both the in-frame small insertion in PDb246 and the frameshifting deletion at the C-terminus might perturb this scaffolding function without eliminating the protein. For the *porU-porV* operon (*porUV*) locus, PDb242 carries a IS982 family transposable element (Fig S2C) insertion (1,716 bp) located 3 bp upstream of the *porU* start codon, a position predicted to abolish transcription or translation initiation of the *porUV* operon (Fig 4D). PorU is a gingipain-fold outer membrane enzyme that cleaves the conserved C-terminal secretion signal from T9SS cargo substrates and their covalent attachment to the cell surface [46,48,49], while PorV is a periplasm-facing lipoprotein that recognizes and shuttles substrates from the SprA pore to PorU for processing [46,51]; loss of both proteins is therefore predicted to block the terminal steps of T9SS-dependent surface delivery. PDb256 carries a complex 33 kb duplication and insertion at the *gldJ* locus (Fig 4D). GldJ is an outer membrane-associated lipoprotein that is essential for stable assembly of the T9SS/gliding motility complex, and its absence leads to destabilization of multiple other complex components including GldK, GldL, GldM, and GldN [46,55,68]. Finally, PDb257 carries an A-to-T transversion in *gldK* introducing a premature stop codon at position K428, predicted to truncate the C-terminus of GldK outer membrane lipoprotein and disrupt its role in anchoring the T9SS complex to the outer membrane and maintaining the structural integrity of the secretion apparatus [46,53] (Fig 4E).

All of the T9SS mutants exhibited a consistent and substantial delay in *C. elegans* killing kinetics relative to wild-type PDb101, with mean survival extended from approximately 25–45 hours on wild-type bacteria (PDb101) to 120–240 hours on the mutant strains (Fig 4F–I, see Supplementary Table S4). To exclude the possibility that the virulence attenuation of T9SS/gliding motility mutants reflects a general growth defect or delay, we measured the growth kinetics of all mutant strains in complex media liquid culture. All T9SS/gliding motility mutants reached similar exponential growth rates and final optical densities as wild-type PDb101, with the exception of a marginal decrease in PDb256 (Fig S6A-H), demonstrating that the virulence attenuation observed in killing assays is not attributable to a generic impairment or delay in bacterial growth.

**Fig S6.**
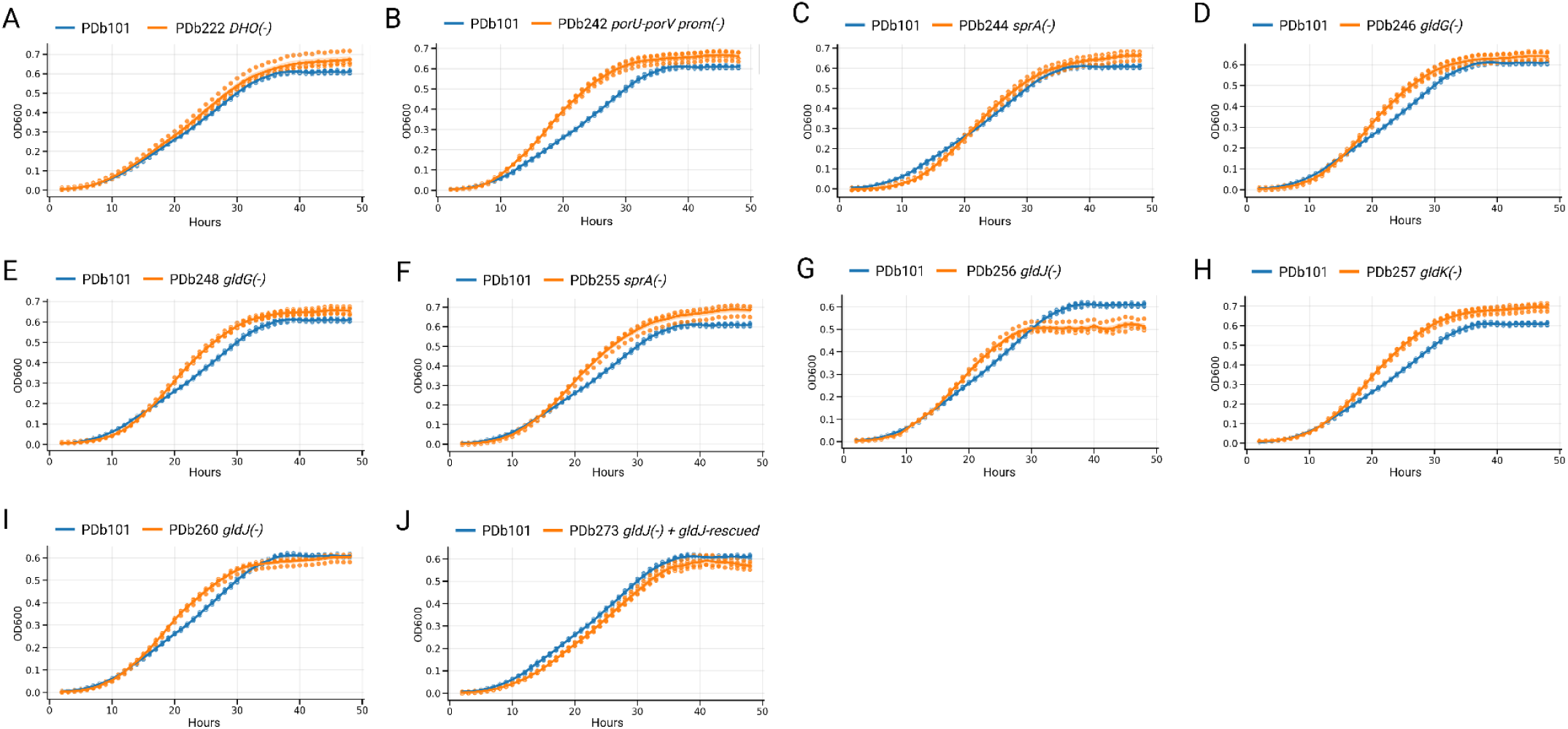
Mutations in the T9SS/gliding motility complex have marginal effects on growth in complex media. (**A-J**) OD600-based growth curves of wild-type PDb101 (blue) and the indicated mutant strains (orange) measured for 48 hours in liquid 1X PGM medium. Each panel compares wild-type PDb101 to a single mutant strain: (**A**) PDb222 *DHO(-)*, (**B**) PDb242 *porU-porV promoter(-)*, (**C**) PDb244 *sprA(-)*, (**D**) PDb246 *gldG(-)*, (**E**) PDb248 *gldG(-)*, (**F**) PDb255 *sprA(-)*, (**G**) PDb256 *gldJ(-)*, (**H**) PDb257 *gldK(-)*, (**I**) PDb260 *gldJ(-)*, and (**J**) PDb273 *gldJ(-)*; *gldJ(+)*-rescued. Lines represent the mean of four biological replicates; shaded areas indicate standard error of the mean (SEM). The wild-type PDb101 growth curve shown in each panel represents the same biological replicate experiment and is reproduced across all panels for direct comparison. All T9SS/gliding motility mutants and the *DHO(-)* auxotrophic control strain reach similar final optical densities as wild-type PDb101 except PDb256 (G) with marginal decrease, indicating that virulence attenuation in these strains is not attributable to a general growth defect in complex medium.

### Genetic complementation and targeted mutagenesis confirm the requirement of T9SS for virulence

To assess whether the virulence attenuation was specifically attributable to T9SS loss rather than to possibly-undetected secondary lesions elsewhere in the genome, we performed genetic complementation analysis. We re-introduced wild-type *gldJ* allele (Fig 5A) into *gldJ(-)* PDb256 background (Fig 5B) using the targeted insertion of plasmid containing a single homology arm designed to recover the truncated *gldJ* locus back to the intact *gldJ* (PDb273, Fig 5C). The re-introduction of a wild-type *gldJ* allele into the *gldJ(-)* mutant PDb256 restored *C. elegans* killing activity to near wild-type levels (Fig 5E), demonstrating that loss of *gldJ* function is the causative lesion in this strain. Furthermore, using the targeted plasmid insertion method (Fig S5B), we disrupted the wild-type *gldJ* locus (PDb260, Fig 5D). The independently generated targeted insertion allele of *gldJ* in PDb260 recapitulated the virulence attenuation phenotype of PDb256 (Fig 5F), providing unambiguous genetic evidence that *gldJ* disruption alone is sufficient to compromise nematode-killing activity and excluding the possibility that the attenuated phenotype of PDb256 reflects unintended consequences of its associated 33 kb duplication. PDb260 also showed similar growth kinetics to that of wild-type PDb101 (Fig S6I), indicating that the reduced virulence by targeted mutagenesis of *gldJ* is not due to bacterial growth defect.

**Fig 5.**
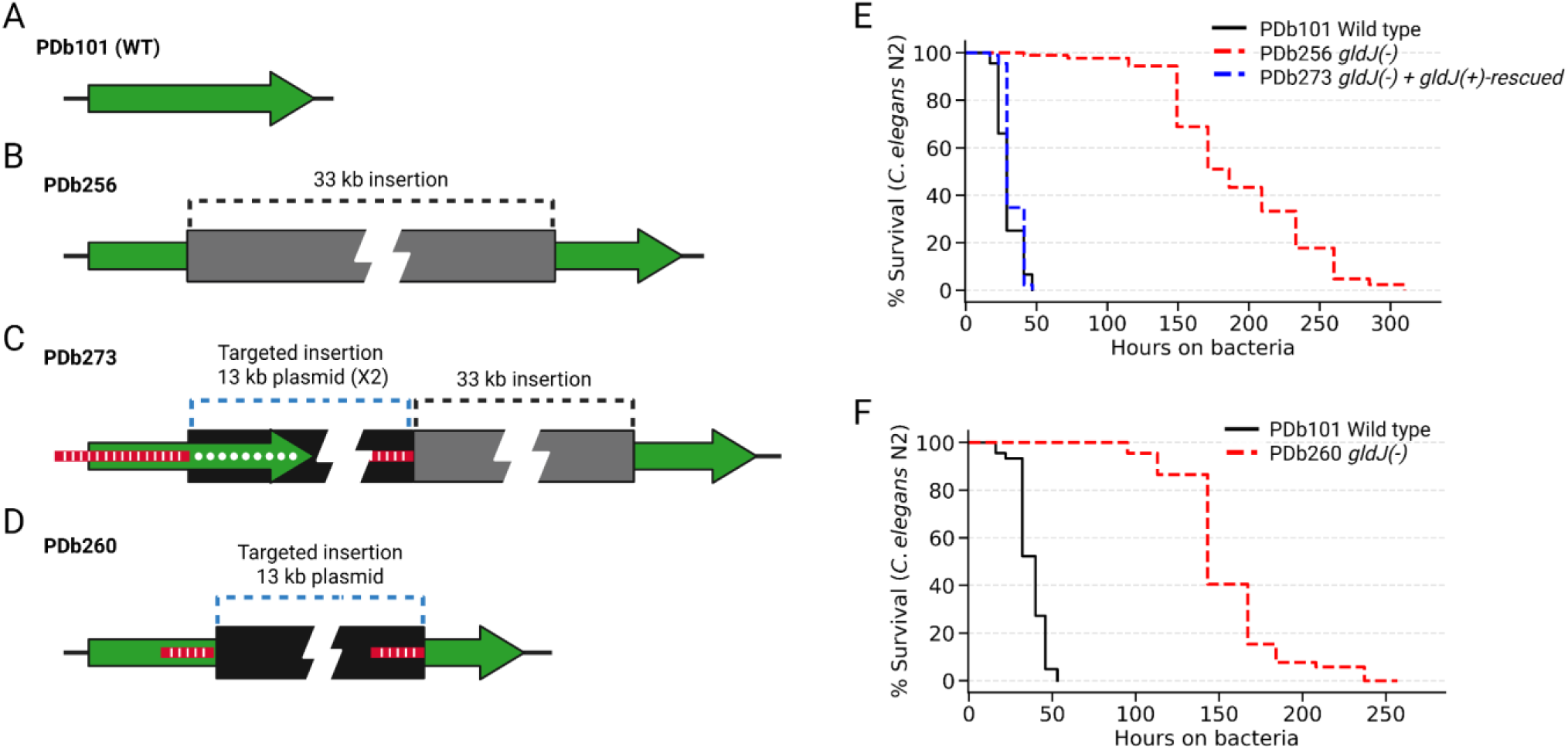
Complementation and targeted mutagenesis of *gldJ* locus confirm the role of *gldJ* in virulence. **(A–D)** Schematic representations of the *gldJ* locus in wild-type PDb101 (A) and mutant strains (B-D). The wild-type *gldJ* locus in PDb101 (WT), shown as a single intact gene (green arrow) (A). PDb256 carries a 33 kb duplication and insertion at the *gldJ* locus, resulting in a split, non-functional copy of *gldJ* flanking the inserted sequence (gray box) (B). PDb273 was generated by targeted plasmid insertion into the PDb256 background: a 13 kb plasmid carrying a wild-type *gldJ* rescue construct and a recoded-*DHO* selectable marker (blue dashed black box, insertion of two copies) was introduced upstream of the 33 kb insertion, restoring two functional *gldJ* copies (the part of green arrow with white dots indicates the recovered region) in the chromosomal configuration. Red/white hatched segments indicate the homology arm regions used for single-crossover recombination. (**E–F**) Kaplan-Meier survival curves of *C. elegans* N2 animals fed on wild-type PDb101 (black) or the indicated mutant strains (red or blue dashed). (**E**) A *gldJ(-)* allele (PDb256) (B) shows virulence attenuation and a *gldJ(-)* strain carrying wild-type *gldJ* rescue construct (PDb273) (C) shows comparable virulence to that of wild-type PDb101, demonstrating almost complete restoration of killing activity upon *gldJ* complementation. (**F**) A *gldJ(-)* allele independently generated by targeted mutagenesis (PDb260) (D) shows substantial attenuation of the virulence, confirming the virulence attenuation phenotype is due to a defect in *gldJ* locus. See Supplementary Table S4 for additional repeats and statistical analysis for the survival data.

### A diverse but functionally biased PDb101 T9SS secretome

To systematically identify the repertoire of proteins secreted by the PDb101 T9SS, we searched for the presence of the conserved C-terminal domains characteristic of T9SS substrates in numerous studied systems [43–46], looking across all predicted PDb101 proteins (Fig 6A; see Materials and Methods for details). This search identified 84 candidate T9SS substrates in the PDb101 genome (Supplementary Table S5), representing a diverse but functionally biased secretome. Sequence- and 3-D structure-based functional annotations of the 84 predicted substrates revealed a striking enrichment of proteolytic enzymes (Fig 6B). The two largest categories were peptidases, with 27 proteins (32.14%) carrying predicted protease domains, and proteins of unknown function (27 proteins, 32.14%). The remaining substrates included grappling hook protein-like proteins, collagenase-like proteins, motility adhesins, glycoside hydrolases, nucleases, and other annotated functions. The dominant representation of peptidases among predicted T9SS cargo proteins suggested that extracellular proteolytic activity represents a major biochemical output of the PDb101 T9SS secretome.

**Fig 6.**
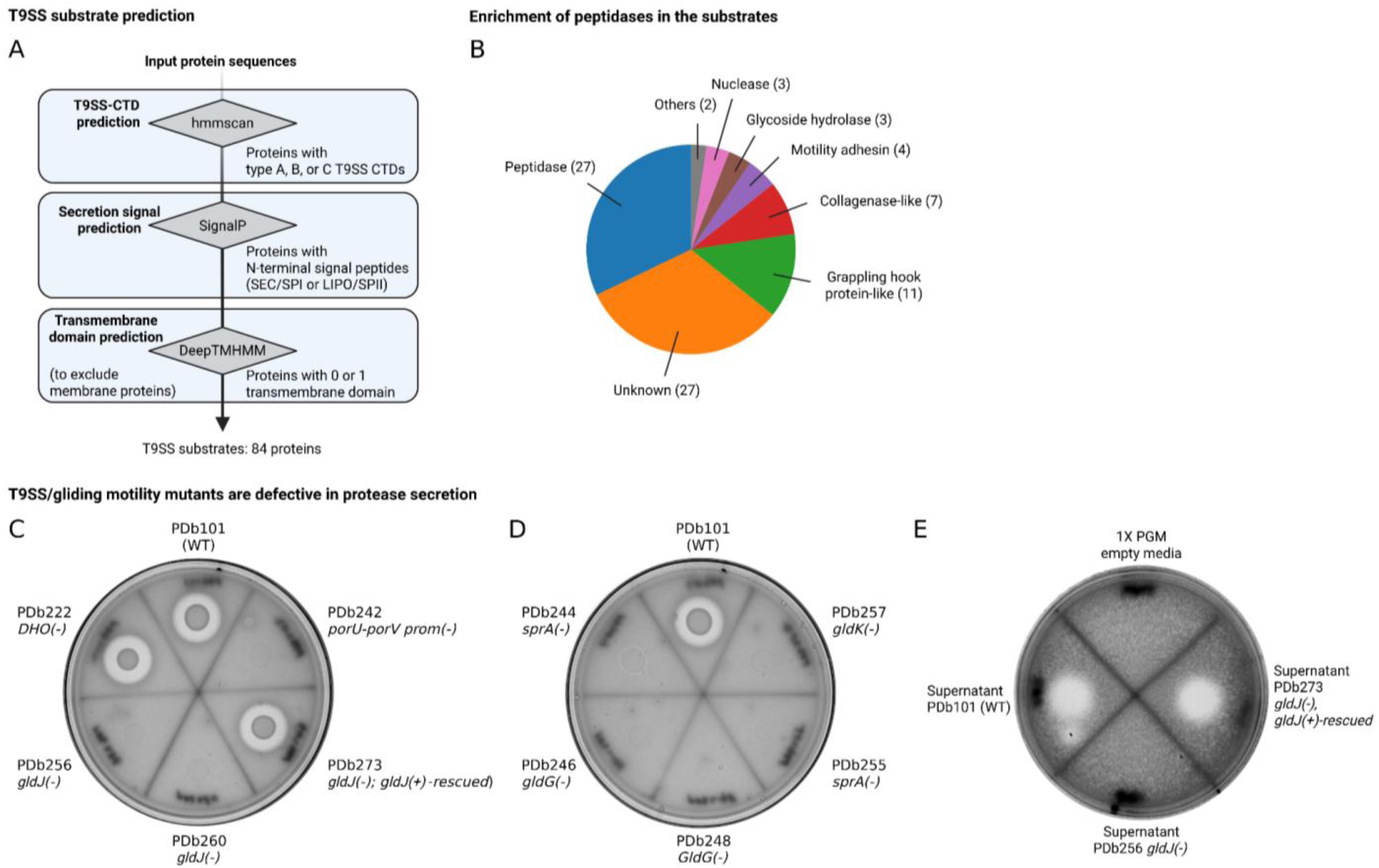
Bioinformatic prediction of T9SS substrates in PDb101 and T9SS-dependent protease secretion. (**A**) Computational pipeline for genome-wide prediction of T9SS cargo substrates in PDb101. Input protein sequences were filtered through three sequential steps: (1) hmmscan-based detection of type A or type B T9SS C-terminal domains (CTDs) using TIGRFAM/Pfam HMM profiles (TIGR04183 and PF18962 for type A, TIGR04131 for type B, and NF033708 for type C); (2) SignalP-based prediction of N-terminal Sec/SPI signal peptides, retaining only proteins with canonical secretion signals; and (3) DeepTMHMM-based transmembrane domain prediction to exclude integral membrane proteins, retaining only proteins with zero or one predicted transmembrane domain. This pipeline identified a total of 84 candidate T9SS substrates in the PDb101 genome. (**B**) Functional classification of the 84 predicted T9SS substrates. The largest categories are peptidases (27 proteins, 32.14%) and proteins of unknown function (27 proteins, 32.14%), followed by grappling hook protein-like proteins, collagenase-like proteins, motility adhesins, glycoside hydrolases, nucleases and other annotated functions. The numbers in parenthesis indicate protein count. (**C–D**) Protease secretion activity of wild-type PDb101 and T9SS/gliding motility mutants assessed by skim milk clearing assay. Strains were spotted onto minimal media agar plates containing skim milk and incubated to allow protease secretion; a clear halo surrounding a colony indicates extracellular protease activity. Wild-type PDb101 and the *DHO(-)* auxotrophic control strain PDb222 produce prominent clearing halos, whereas the *porU-porV promoter(-)* mutant PDb242, both *gldJ(-)* mutants (PDb256, PDb260) show markedly reduced or absent clearing (C). Protease secretion is restored in PDb273, consistent with complementation of the *gldJ* lesion (C). *sprA(-)* mutants (PDb244, PDb255), *gldG(-)* mutants (PDb246, PDb248), and the *gldK(-)* mutant (PDb257) all exhibit reduced extracellular protease activity compared to wild-type PDb101 (D). (**E**) Protease activity of culture supernatants assessed on skim milk agar plates. Cell-free culture supernatants from wild-type PDb101, the *gldJ(-)* mutant PDb256, and the *gldJ(-); gldJ(+)-rescued* strain PDb273 were concentrated 100-fold using centrifugal filters and spotted directly onto skim milk agar plates, alongside a 100-fold concentrated 1X PGM empty media as a control. Supernatant from PDb101 and from the *gldJ(-); gldJ(+)-rescued* strain PDb273 produced clear zones of skim milk hydrolysis, whereas supernatant from the *gldJ(-)* mutant PDb256 and the media-only control showed no clearing, confirming that the secretion defect in *gldJ(-)* reflects a loss of extracellular protease activity rather than a failure of bacterial growth on the plate.

### The attenuated virulence of T9SS mutants is accompanied by perturbed secretion of extracellular proteases

To assess whether the T9SS/gliding motility mutants identified in this study are defective in functional protease secretion, we performed skim milk clearing assays, in which clearing of the milk turbidity serves as a readout for extracellular protease competence [69]. Wild-type PDb101 produced prominent clearing halos, as did the *DHO(-)* auxotrophic control strain PDb222, confirming that pyrimidine auxotrophy does not affect T9SS function (Fig 6C). In contrast, all T9SS/gliding motility mutants tested showed markedly reduced skim milk clearing, including the *porU-porV* promoter mutant PDb242, both independent *gldJ(-)* alleles (PDb256 and PDb260), both independent *sprA(-)* alleles (PDb244 and PDb255), both independent *gldG(-)* alleles (PDb246 and PDb248), and the *gldK(-)* mutant PDb257 (Fig 6C-D). The protease secretion defect of the *gldJ(-)* mutant PDb256 was rescued by re-introduction of wild-type *gldJ* allele in PDb273, confirming that the secretion deficiency is specifically attributable to loss of *gldJ* function (Fig 6C). Furthermore, cell-free supernatants of wild-type PDb101 also produced clearing halos whereas an empty media control or the supernatants of *gldJ(-)* mutant PDb256 did not (Fig 6D). Re-introduction of wild-type *gldJ* allele in PDb273 restored halo-forming activity, confirming proteolytic activities of secreted proteases via T9SS (Fig 6D). Taken together, these results demonstrate that all mutant loci identified in this study are required for effective T9SS-dependent extracellular protein secretion in PDb101, establishing that these mutations compromise general secretion of T9SS substrates rather than any single effector.

## Discussion

In this study, we established the first genetic manipulation system for *Chryseobacterium nematophagum* with a pyrimidine auxotrophy-based selection system. By using a single homology arm recombination strategy for conjugation-mediated chromosomal disruption, we enabled targeted genetic manipulation at any locus in the PDb101 genome for the first time. Leveraging this system with forward/reverse genetics, we identified the Type IX secretion system (T9SS) machinery as a central virulence determinant of PDb101 against *C. elegans*. Independent mutations in five T9SS/gliding motility genes each substantially attenuated nematode killing and abolished extracellular protein secretion, and these phenotypes were confirmed through targeted mutagenesis and genetic complementation. Together, these findings establish *C. nematophagum* PDb101 as a genetically tractable model for dissecting bacterial pathogen-host nematode interactions and the T9SS-dependent bacterial virulence, adaptation and evolution mechanisms at the molecular level.

### Ecological and evolutionary characteristics of nematophagous *Chryseobacterium*

The isolation of PDb101 from California soil extends the known geographic distribution of nematophagous *C. nematophagum* from Europe and Asia to North America. It is consistent with the cosmopolitan distribution of its host nematodes across terrestrial ecosystems. The high degree of genomic similarity between PDb101 and geographically distant nematophagous strains JUb129 (France) and JUb275 (India), reflected in greater than 93.5% average nucleotide identity, suggests that the nematophagous lifestyle and its associated genetic determinants are conserved within this species. Nevertheless, PDb101 is distinguished from other *C. nematophagum* isolates by a markedly elevated transposon burden, with IS elements broadly distributed throughout the genome and signatures of past transposon burst activity (Fig 1D-F). This elevated genome plasticity may reflect exposure to distinct mobile genetic element communities in North American soil environments and could contribute to strain-level variation in secretome composition, regulatory architecture, or host range. The substantial diversity of CRISPR spacer arrays across PDb101, JUb129, and JUb275 further supports the interpretation that these geographically isolated populations have been exposed to distinct phage and mobile genetic element communities, consistent with independent evolutionary trajectories following dispersal from a common ancestor. Together, these observations suggest that while nematophagous activity is a conserved species-level trait in *C. nematophagum*, individual strains have undergone considerable genome-level diversification, and that PDb101 may represent a particularly dynamic lineage within this species.

### T9SS as a virulence apparatus to nematodes

The convergence of independent mutations in five T9SS/gliding motility genes on a single virulence-attenuated phenotype establishes the T9SS/gliding motility complex as essential for PDb101 nematophagous activity. The genetic evidence is notably robust: (i) each of five loci was disrupted by independent mutational events, (ii) allelic pairs were recovered for *sprA* and *gldG*, (iii) the causative role of *gldJ* disruption was confirmed by both targeted mutagenesis and complementation analysis, and (iv) the concordant loss of extracellular protein secretion in all mutants confirms that virulence attenuation reflects a general secretion defect rather than the incidental loss of any single effector.

Bacterial protein secretion systems are central mediators of host-pathogen interactions, enabling the delivery of toxins, adhesins, and immune-modulatory effectors directly to host cells or tissues [70,71]. Distinct secretion systems have been linked to virulence across diverse bacterial pathogens. For example, the Type III and Type IV secretion systems inject effector proteins directly into host cells in many Gram-negative pathogens [72–75], while the Type VI secretion system mediates both interbacterial competition and, in some species, direct host cell intoxication [76,77]. Among *Bacteroidota* specifically, the T9SS has emerged as a comparably central virulence apparatus, exporting a structurally diverse cargo of surface-associated and secreted effectors that mediate host tissue damage and immune evasion [44–46,78].

Prior to this study, T9SS-dependent virulence had been characterized exclusively in the context of vertebrate host infection: in the human periodontal pathogens *Porphyromonas gingivalis*, T9SS-dependent secretion of gingipain proteases drives tissue destruction and immune evasion [42,58–63]; in the fish pathogens *Flavobacterium columnare* and *F. psychrophilum*, T9SS-dependent proteases and adhesins are required for virulence in zebrafish and salmonids [64,66,79]; and in the avian pathogen *Riemerella anatipestifer*, T9SS-secreted serine proteases contribute to complement evasion and systemic infection in ducks [80,81]. Together, our findings provide the first demonstration that T9SS machinery constitutes a central virulence determinant in a nematophagous bacterium, establishing that T9SS-dependent pathogenesis is not restricted to vertebrate hosts but extends to invertebrates and expanding the ecological diversity of host-pathogen interactions that T9SS mediates. Furthermore, our findings add to a growing body of evidence that the T9SS represents a remarkably adaptable secretion platform whose cargo composition has been repeatedly co-opted across *Bacteroidota* to suit distinct ecological niches and host interactions. The composition of the PDb101 T9SS secretome and which substrates are responsible for nematophagous activity, therefore represents an important question not only for understanding *C. nematophagum* virulence but also for illuminating how T9SS cargo diversification drives ecological specialization across the *Bacteroidota* phylum.

### Nematophagous effectors within the PDb101 T9SS secretome

Key questions raised by our findings are whether nematophagous activity might be attributable to a single T9SS substrate, and whether the dominant proteolytic component of the secretome represents the essential toxicity effector.

The broad conservation of T9SS machinery and M12B family metalloproteases across non-pathogenic *Chryseobacterium* species argues that any role(s) of these proteases in nematophagous activity would rely either on precise activity of one or a few proteases on nematode-specific targets, on localized release mechanisms targeting nematodes during specific colonization. Likewise, the nematophagous activity could reside among the handful of predicted T9SS substrates with no clear homologs in non-pathogenic *Chryseobacterium* species identified by comparative secretome analysis. Whether any individual *C. nematophagum* T9SS substrate is necessary or sufficient for toxicity remains unresolved, as the experiments reported here inactivate the entire secretion apparatus rather than individual substrates. Systematic individual and combinatorial disruption of *C. nematophagum*-specific T9SS substrate genes will be a valuable next step to identify the effectors responsible for nematophagous activity.

### Mechanistic ambiguity between T9SS-dependent secretion and gliding motility for pathogenicity

A related limitation is that all mutations recovered affect components shared between the T9SS and the gliding motility apparatus, precluding mechanistic separation of the two functions. Every locus disrupted—*sprA*, *gldG*, *gldJ*, *gldK*, and the *porUV* operon—is expected to simultaneously impair T9SS-dependent secretion and, in motile species, surface gliding. The concordant secretion defect in all mutants supports T9SS-dependent secretion as a central virulence mechanism but does not exclude an independent contribution of gliding motility to nematode killing.

In other *Bacteroidota* pathogens, the two activities make distinct contributions to virulence: in *Flavobacterium columnare*, gliding and secretion mutants exhibit partially non-overlapping virulence defects in fish infection models, suggesting that the two activities serve non-redundant roles during pathogenesis [79], while in *Porphyromonas gingivalis*, which lacks gliding, T9SS-dependent secretion alone is sufficient for pathogenesis [42,82]. Although two species in the *Chryseobacterium* genus (*C. gleum* and *C. sp.PMSZPI*) were shown to move through gliding on soft agar surface [83,84], whether PDb101 occupies a similar motile niche remains unresolved. Gliding of PDb101 was not detected under standard or low-nutrient agar plates across a range of agar concentrations for variable surface stiffness (data not shown), yet PDb101 encodes protein homologs of the C-terminal domains of the *F. johnsoniae* motility adhesin RemA, raising the possibility that gliding occurs under conditions not yet tested. Resolving this ambiguity will require both systematic exploration of alternative gliding conditions and the development of separation-of-function alleles to directly dissect their respective contributions to nematophagous activity.

### T9SS-biased mutagenic and selection effect during conjugation

The recovery of multiple independent T9SS/gliding motility mutants from a single conjugation experiment represents a serendipitous observation: despite numerous subsequent conjugation experiments performed under nominally identical conditions, this clustering of T9SS-deficient mutants was never observed again. In *Flavobacterium* species, T9SS and gliding motility mutants are known to form more compact, non-spreading colonies on agar surfaces [42,46]; an analogous colony behavior in PDb101 may have conferred a selective advantage under the specific plating conditions of that batch, providing the most parsimonious explanation for their enrichment. Regardless of the underlying cause, the biological significance of the finding is clear–the independent occurrence of loss- or reduction-of-function mutations across five distinct T9SS/gliding motility loci, each associated with consistent virulence attenuation, provides robust genetic evidence that intact T9SS function is required for PDb101 pathogenicity. This conclusion resulted from independent experimental validation through targeted mutagenesis and complementation analysis rather than on the circumstances of the original recovery of the mutants.

### Limitations and future directions of genetic manipulations in *C. nematophagum* (PDb101)

The auxotrophy-based genetic manipulation system established here represents the first genetic platform for *C. nematophagum* and its close relatives, overcoming the fundamental barrier of broad intrinsic antibiotic resistance that has precluded molecular dissection of this organism. The ability to deliver DNA by conjugation and achieve targeted chromosomal disruption at any locus now makes it possible to systematically interrogate the genetic basis of nematophagous activity in an organism that was previously inaccessible to experimental genetics.

The current system has clear limitations; it relies on a single selectable marker, depends on conjugation for DNA delivery, and is restricted to low-throughput single-crossover insertions that preclude scarless editing. However, these are well-defined technical challenges with tractable solutions established in related *Bacteroidota*, including expansion of auxotrophic markers, electroporation-based transformation, counterselection strategies for allele replacement, and high-throughput transposon random mutagenesis. Looking forward, the endogenous Type II CRISPR-Cas9 system encoded by PDb101 raises the intriguing possibility that the native Cas9 machinery could be co-opted for precision genome editing by supplying appropriate guide RNAs, potentially enabling single-nucleotide editing, CRISPRi-based transcriptional modulation, and high-throughput functional screens without the need for selectable markers at every target locus [85].

Together, the genetic foundation established here and its natural extensions position *C. nematophagum* as an experimentally tractable model for systematic dissection of *Bacteroidota* for virulence mechanisms, evolution for environmental adaptation and bacterium-nematode interactions at the molecular level.

## Materials and Methods

All materials and resources are listed in the Supplementary Table S1.

### *C. elegans* strain and maintenance

*C. elegans* (N2, CGC1) was maintained at 20°C on nematode growth medium (NGM) plates seeded with *E. coli* OP50, as previously described [86].

### Isolation and culture of *C. nematophagum* (PDb101)

Environmental samples, including soil, were collected from several locations in Palo Alto, California, USA, and brought to the laboratory to screen for nematodes and nematode-associated microorganisms. As part of this screen, A *Polintovirus* reporter *C. elegans* strain (PD3732) was transferred into environmental samples to detect nematode-infecting dsDNA virus (*Polintovirus*) [87] activity via reporter signals. During this screen, unexpected nematode killing activity was observed in a sample. To identify the causative organism, bacteria and fungi forming discrete colonies on solid NGM plates were isolated and cultured individually in 2X TY (tryptone/yeast extract) media for 48-72 hours at room temperature. The bacterial and fungal cultures were seeded onto NGM agar plates, and wild-type *C. elegans* (N2, CGC1) were transferred to assess killing activity. A bacterial isolate designated PDb101 was identified based on its reproducible nematophagous activity against wild-type *C. elegans*. We noted that peptone is a better nutrient resource for PDb101 growth than tryptone/yeast extract since a custom formulation designated PDb101 growth medium (1X PGM; 0.25% peptone and 0.3% NaCl) consistently supported faster growth of PDb101 relative to 2X TY medium at 23-27℃. PGM was therefore mainly used for subsequent cultures of PDb101 in this study, otherwise noted.

### *C. elegans* survival assays

For survival assays, single colonies of OP50 or PDb101 (wild-type or mutant) strains were cultured overnight in 2X TY medium (37°C) and for 48 hours in 1X PGM (23°C), respectively. Liquid cultures were seeded onto NGM plates and incubated overnight prior to use. *C. elegans* were grown on OP50-seeded plates at 20°C until the young adult stage, then transferred (90 animals per condition; 30 animals per plate) to plates seeded with OP50, wild-type PDb101, or PDb101 mutant strains, and maintained at 20°C throughout the assay. Worms were transferred daily to fresh plates until all animals had died or ceased egg-laying. Animals were scored two to three times daily during the first two-three days and once daily thereafter; animals that failed to respond to gentle prodding or exhibited internal hatching (natural impact by PDb101) were scored as dead. Animals that displayed intestinal extrusion, crawled onto the wall of the plate, or burrowed into the agar and could not be reliably tracked were excluded from the dead count and recorded as censored; censored animals were nonetheless included in the statistical analysis. Statistical analyses were performed using OASIS 2 [88], and *p*-values were calculated by the log-rank test. For heat-killing of PDb101, the saturated PDb101 cells were concentrated 10 times by centrifugation at 1,500 x *g* and resuspension of the cells. The concentrated cells were fully submerged in a 70°C water bath from the top lid to the bottom for 25 minutes to ensure complete and uniform heat inactivation of the bacterial cells. Heat-killed PDb101 cells were then seeded onto NGM plates and incubated overnight at room temperature prior to worm transfer. *C. elegans* animals were transferred to heat-killed PDb101-seeded plates and survival was assessed as described above. Where indicated, 5-fluoro-2′-deoxyuridine (FUdR) was applied to PDb101-seeded plates 24 hours prior to worm transfer to kill PDb101. Briefly, 120 µL of 100X FUdR stock solutions (5 µM, 50 µM, 500 µM, or 5 mM) were treated directly onto the bacterial lawn and allowed to absorb, yielding final plate concentrations of 0.05 µM, 0.5 µM, 5 µM, or 50 µM FUdR respectively in 12 mL NGM plates.

### Bacterial genome sequencing

High molecular weight (HMW) genomic DNA was extracted from liquid cultures of PDb101 using the Zymo Quick-DNA HMW MagBead Kit according to the manufacturer’s instructions. Long-read whole-genome sequencing was performed through Plasmidsaurus’ bacterial whole-genome sequencing platform. Briefly, the sequencing library was prepared using an amplification-free long-read protocol with Oxford Nanopore v14 library prep chemistry, including minimal, sequence-independent fragmentation of input gDNA via tagmentation. The library was sequenced on R10.4.1 flow cells. Raw reads were basecalled using Dorado (https://github.com/nanoporetech/dorado) with the v4.3 Super-Accurate model and a default Q10 quality filter. *de novo* assembly was performed using Autocycler [89] with three assemblers–Flye [90,91] v2.9.6, Hifiasm [92], and Plassembler [93] v1.8.0–followed by consensus resolution and contig rotation using dnaapler [94]. For short-read sequencing, a library was constructed from the same HMW gDNA using the Nextera XT kit with tagmentation and 12 cycles of PCR amplification and sequenced on the Illumina MiSeq platform. Short reads were aligned to the long-read assembly using BWA-MEM [95] and used to polish the assembly. Variants were called using BCFtools [96] and corrected in the long-read assembly. Genome annotation was performed with Bakta [97] v1.11, and genome completeness and contamination were assessed with CheckM [98] v1.2.2. Genomic DNA from additional PDb101 mutant strains was also subjected to whole-genome sequencing through either the Illumina MiSeq short read sequencing platform or the Plasmidsaurus ONT long-read sequencing platform. Raw reads were aligned to the completed PDb101 reference genome using BWA-MEM [95] or minimap2 [99,100], and variants were called from the resulting alignments with BCFtools^102^ to identify mutations in each strain.

### Chryseobacterium comparative genomic analyses

Genome assemblies and 16S rRNA gene sequences from selected *Chryseobacterium* species and *Riemerella anatipestifer* were downloaded from the NCBI Genome Assembly and Nucleotide databases (listed in the Supplementary Table S2). For phylogenetic analysis, genome sequences were annotated with Prokka [101] and compared using Roary [102] to generate a core genome alignment. A maximum-likelihood phylogenetic tree was constructed from the resulting multiple sequence alignment using IQ-TREE [103–105]. The tree was rooted with *R. anatipestifer* as the outgroup and visualized using the Python ETE3 package [106]. Average nucleotide identity (ANI) between genome sequences and between 16S rRNA gene sequences was calculated using pyANI-plus [107] with the MUMmer alignment mode. CRISPR array sequences in three *C. nematophagum* genome assemblies were identified from Bakta [97] annotations, and individual repeat and spacer sequences were verified and compared by manual inspection. Insertion sequence (IS) elements were predicted using ISEScan [108]. Only elements with an intact transposase coding sequence and detectable terminal inverted repeats (TIRs) were retained and included in figures. Genomic features–including CDS, rRNA, tRNA, CRISPR arrays, and insertion sequences–were visualized for each genome using Circos [109] with pyCirclize (https://github.com/moshi4/pyCirclize).

### PDb101 EMS mutagenesis

To optimize conditions for ethyl methanesulfonate (EMS) mutagenesis, multiple combinations of EMS dose, incubation time, and bacterial growth phase were evaluated (Supplementary Table S3). For the optimized protocol, a single colony of PDb101 was inoculated into 1X PGM and incubated with gentle shaking for 48 hours at 23°C to reach saturation phase. The saturated culture was then diluted 1:100 into 2X TY medium and incubated for 16 hours at 23°C to obtain early log-phase cells. Cells were washed once by centrifugation at 1,500 x *g* and resuspension in fresh 2X TY medium. EMS was added to a final concentration of 1% (v/v), and cells were incubated with gentle shaking for 4 hours at room temperature (RT). EMS-treated cells were washed three times with fresh 2X TY medium and then allowed to recover with gentle shaking for 4 hours at RT. Recovered cells were serially diluted and plated onto NGM plates to obtain single colonies. To confirm EMS activity in each mutagenesis experiment, a portion of the treated culture was plated onto NGM plates containing rifampicin (0.1 µg/mL) to detect EMS-induced rifampicin-resistant mutants as a positive indicator of mutagenesis. For mutant screening, individual colonies (over 4,000 colonies) were picked into 1X PGM in 96-well plates and incubated for 48 hours. Cultures were then diluted 1:100 into NM9GTA minimal medium (see below), and strains failing to grow were identified as auxotrophic mutants. This screen identified a dihydroorotase (*DHO*)-deficient mutant (PDb222) and a carbamoyl phosphate synthase (CPS) large subunit-deficient mutant (PDb223).

### Chemically defined minimal media for PDb101 growth

A nearly minimal medium, designated NM9GTA (N = nearly minimal, M9 = M9 salts base, G = glucose, T = thiamine, A = amino acids), was formulated to support the growth of *C. nematophagum* PDb101 under defined nutritional conditions. 1X NM9GTA contains the following components: 0.6% Na_2_HPO_4_, 0.3% KH_2_PO_4_, 0.05% NaCl, 0.1% NH_4_Cl, 1 mM MgSO_4_, 0.1 mM CaCl_2_, 0.1% glucose, 50 µg/L thiamine hydrochloride, 1X MEM Non-Essential Amino Acids Solution (without L-glutamine), and 1X MEM Amino Acids Solution (without L-glutamine). For solid medium, a 2X NM9GTA solution was prepared and mixed in equal volumes with autoclaved 3% agar, yielding a final medium of 1X NM9GTA in 1.5% agar. Where required, uracil was supplemented to a final concentration of 50 µg/mL (NM9GTA-uracil) to support the growth of mutant PDb101 cells defective in *de novo* pyrimidine biosynthesis auxotrophic strains. If required, ampicillin (75-100 µg/mL) was supplemented to 1X NM9GTA to select PDb101 by killing ampicillin-sensitive *E. coli*.

### Multiple sequence alignment

Selected bacterial and eukaryotic dihydroorotase (DHO, type I) and carbamoyl phosphate synthetase (CPS) protein sequences were downloaded from the NCBI Protein database. In bacteria, DHO (type I) and CPS are encoded as separate polypeptides, whereas in eukaryotes both activities are carried by a single multifunctional CAD protein (CPS-ATCase-DHO). For alignment, the DHO and CPS domains were extracted from eukaryotic CAD sequences and analyzed separately alongside their bacterial counterparts. Sequences were aligned using MAFFT [110] with the G-INS-i option, and low-information-content positions were removed using trimAl [111] with the gappyout option. The trimmed alignment was visualized in ClustalX [112] using the default ClustalX color scheme.

### Cloning and plasmid construction

To construct a broad-host-range plasmid carrying the wild-type *DHO* locus of PDb101, the plasmid pBroad-T7RNAP-chlorR (Addgene #199992) [113] was digested with NruI and StuI to isolate the broad-host-range backbone carrying a chloramphenicol-resistance cassette. The PDb101 *DHO* element (comprising the promoter, coding sequence, and 3′ downstream region) was amplified by PCR using PDb101 genomic DNA as template and assembled into the linearized backbone by Gibson assembly to generate pDEJ125. To construct *DHO* complementation plasmids carrying heterologous *DHO* alleles, the *DHO* coding sequence from JUb44 (*Chryseobacterium scophthalmum*) was amplified by PCR from JUb44 genomic DNA, and a codon-recoded *DHO* sequence (re-coded-DHO) was synthesized as a double-stranded gBlock (Integrated DNA Technologies). Each insert was Gibson-assembled into the linearized broad-host-range backbone to generate pDEJ160 (JUb44-DHO) and pDEJ167 (re-coded-DHO), respectively. For chromosomal disruption of *gldJ*, a 507 bp internal fragment of the *gldJ* coding sequence was amplified from PDb101 genomic DNA and Gibson-assembled with the linearized broad-host-range backbone carrying the re-coded*-*DHO selection cassette to generate pDEJ156. Upon conjugation into PDb101, plasmid insertion via single-crossover recombination at the internal *gldJ* fragment disrupts the chromosomal *gldJ* locus (see the section below). For restoration of wild-type *gldJ* in *gldJ* mutants (PDb256), a construct carrying the intact *gldJ* coding sequence flanked by 1,729 bp of upstream and 239 bp of downstream sequence was amplified as a single PCR product from PDb101 genomic DNA and Gibson-assembled with the linearized broad-host-range backbone carrying the re-coded-DHO selection cassette to generate pDEJ174. All the assembled plasmids were transformed into the TOP10 chemically competent cells, and the transformant cells were selected on 2X TY media containing chloramphenicol (34 µg/mL). All the plasmids were amplified and extracted from the transformant cells by using NucleoSpin-Plasmid kit (Takara Bio USA, Inc.), and sequence-verified from sequencing using the Plasmidsaurus plasmid sequencing platform.

### Conjugation-Mediated Plasmid Delivery and *DHO* complementation

For conjugation-mediated plasmid delivery into PDb101, the *E. coli* strain BW19851 [114] was used as the donor and the *DHO*-deficient PDb101 mutant (PDb222) was used as the recipient. Chemically competent BW19851 cells were prepared as described in Chung et al. (1989, 1993) [115,116]. Each plasmid (pDEJ125, pDEJ160, or pDEJ167) was transformed into competent BW19851 cells, and transformants were selected on 2X TY agar (1.5%) media containing chloramphenicol (34 µg/mL). A single transformant colony was cultured overnight at 37°C in liquid 2X TY medium, then diluted 1:100– 1:200 into fresh 2X TY medium and incubated for 2.5 hours to obtain mid-log phase donor cells. In parallel, a single colony of PDb222 was inoculated into 1X PGM and cultured for 48 hours at 23°C, then diluted 1:100 into 2X TY medium and incubated for 16 hours to obtain early log-phase recipient cells. Mid-log phase BW19851 and early log-phase PDb222 cells were each collected and concentrated 50 times by centrifugation at 1,500 × *g* at room temperature, resuspended in 15–25 µL of 2X TY medium, and combined. The resulting 30–50 µL cell mixture was deposited onto a 0.45 µm mixed cellulose ester (MCE) membrane filter (MF-Millipore, 24 mm) placed on an NGM agar plate (60 mm dish) and incubated at room temperature for 24 hours to allow mating. Following conjugation, cells were recovered by washing the membrane filter with 4 mL of 1X phosphate-buffered saline (PBS). The cell suspension was concentrated 10-fold by centrifugation at 1,500 x *g* for 5 minutes and plated onto NM9GTA minimal agar supplemented with ampicillin (75–100 µg/mL) to counter-select against residual BW19851 donor cells. Plates were incubated at 27°C for 48–72 hours. Successful *DHO* complementation was confirmed by the appearance of PDb222 transconjugant colonies, whose morphology and pigmentation distinct from *E. coli* BW19851 colonies and thus readily distinguishable on plate. Subsequently, 6–12 transconjugant colonies were picked and restreaked onto fresh NM9GTA-Amp minimal agar plates and incubated for an additional 48–72 hours to confirm the absence of BW19851 contamination. The transconjugant colonies then were cultured in liquid NM9GTA or 1X PGM for 48 hours (23°C) to extract HMW genomic DNA and the sequences were verified by whole-genome sequencing through the Plasmidsaurus bacterial whole-genome sequencing platform.

### Targeted mutagenesis and complementation of the *gldJ* locus

For targeted disruption of *gldJ*, pDEJ156 was delivered into the *DHO*-deficient strain PDb222 by bacterial conjugation with *E. coli* BW19851 as described above. Plasmid insertion via single-crossover recombination at the internal *gldJ* fragment disrupts the chromosomal *gldJ* locus. Transformants were screened by PCR genotyping and sequence-verified by long-read sequencing through the Plasmidsaurus bacterial whole-genome sequencing platform. For rescue of the *gldJ* disruption mutant, pDEJ174 was introduced into PDb256–a *gldJ* disruption mutant maintaining a *DHO*-deficient genetic background–by the same conjugation procedure. Plasmid insertion via single-crossover recombination at the *gldJ* locus restored a wild-type copy of *gldJ*. The extended upstream homology arm (1,729 bp) in pDEJ174 was included to favor recombination at the correct locus. Transformants with plasmid insertions were screened by PCR genotyping and sequence-verified as well by long-read sequencing through the Plasmidsaurus bacterial whole-genome sequencing platform.

### *in silico* prediction of type IX secretion system (T9SS) substrates

T9SS substrates among predicted PDb101 proteins were identified using a three-step computational pipeline. First, conserved C-terminal domain (CTD) sequences were detected by hmmscan [117] using profile HMMs for Type A CTDs (TIGR04183.1 and PF18962) and Type B CTDs (TIGR04131.1). Only proteins with a significant CTD match at the C-terminus were retained. Second, the presence of an N-terminal signal peptide was assessed using SignalP [118]; proteins were retained if they carried a Sec/SPI or Lipo/SPII signal peptide with a prediction probability ≥ 0.5. Third, transmembrane domain content was evaluated using DeepTMHMM [119], and proteins predicted to contain more than two transmembrane domains were excluded to remove putative integral membrane proteins from the candidate list. Proteins passing all three filters were designated predicted T9SS substrates. A total of 84 proteins were identified as predicted T9SS substrates in PDb101, and subjected to functional annotation by HHpred [120] homology search and/or AlphaFold3 [121] structural prediction followed by Foldseek [122] structural homology search.

### Skim milk plate extracellular protease assay

To assess extracellular protease activity dependent on the T9SS, a skim milk agar medium was prepared with the following composition: 0.6% Na_2_HPO_4_, 0.3% KH_2_PO_4_, 1 mM MgSO_4_, 0.1 mM CaCl_2_, 50 µg/mL thiamine hydrochloride, 50 µg/mL uracil, 1.5% skim milk, and 1.5% agar. Briefly, 500 mL of 2X M9 base solution (1.2% Na_2_HPO_4_ and 0.6% KH_2_PO_4_) and 500 mL of 3% agar solution were separately autoclaved. Skim milk (15 g) was added directly to the hot 2X M9 base solution for heat sterilization. After cooling to 55°C, MgSO_4_, CaCl_2_, thiamine hydrochloride, and uracil, all of which were filter-sterilized, were added. Then the supplemented M9 base solution was mixed with the autoclaved 3% agar solution to yield the final 1X medium. Carbon and nitrogen sources (glucose, NH_4_Cl, and amino acids) were intentionally omitted to compel bacteria to rely exclusively on extracellular proteolysis of skim milk proteins as the sole nutrient source, such that growth on this medium is strictly dependent on extracellular protease secretion and activity. Wild-type PDb101 and all mutant strains were cultured in 1X PGM for 48 hours, and cell densities were normalized by optical density at 600 nm (OD600). OD600-normalized cell suspensions (5 µL) were spotted onto skim milk agar plates and incubated at 27°C for 48–72 hours. Plates were imaged using a BioRad GelDoc Go Imaging System.

## Acknowledgements

We are thankful for the financial support of NIH (Grant GMR35130366 to AZF), the Human Frontier Science Program (Long-Term Postdoctoral fellowship LT000329/2019-L to DEJ), the Bernard Cohen Postdoctoral Research Fellowship (to DEJ). We are grateful to our colleagues for their support and suggestions during this work: K. Artiles, O. Ilbay, U. Enam, D. Galls, J. Kim, C, Benko, H. Krupkin, D. Lipman, L. Wahba, M. McCoy, M. Shoura, N. Hall, I. Zheludev, N. Jain, S. Sundani, S. Bowden, G. Heo, Y Kim, A. Page, M. Felix, K. Perez, M. Guzman, and M. Kwon. We acknowledge suggestions in sentence construction, translation and reference gathering from Claude (Anthropic); all of these suggestions were assessed sentence-by-sentence and word-by-word, and adopted, discarded, or corrected as appropriate.

## Data availability

The PDb101 genome assembly and raw whole-genome sequencing reads have been deposited in NCBI Genome and the Sequence Read Archive (SRA) database, respectively, under BioProject accession PRJNA1512035.

